# Labeling proteins within *Drosophila* embryos by combining FRET reporters, position-specific genomic integration, and GAL4-reponsive expression

**DOI:** 10.1101/743492

**Authors:** Tzyy-Chyn Deng, Chia-Jung Hsieh, Michael De Freitas, Maria Boulina, Nima Sharifai, Hasitha Samarajeewa, Tatsumi Yanaba, James D. Baker, Michael D. Kim, Susan Zusman, Kenneth H. Wan, Charles Yu, Susan E. Celniker, Akira Chiba

## Abstract

Protein interaction network (PIN) or interactome has been mapped vigorously for the entire genome. We recognize, nonetheless, that such a map could illuminate profound insights had its context been revealed. We describe a scalable protein lableling method that could re-supply natural context back to the map of protein interactome. Genetically encoded fluorescent proteins, position-specific genomic integration and GAL4-responsive expression control enable labeling proteins *A, B* and *C* each with a either an eGFP, mCherry or NirFP in specified cells of optically transparent animals such as *Drosophila* embryos. While following multiple proteins through development and behavior, these labels offer separable pairs of Förster resonance energy transfer between proteins *A* and *B* and proteins *B* and *C*. We test and observe FRET interactions between specific protein pairs controlling cytoskeleton, nuclear signaling and cell polarity. By using our protein labeling method, it will be possible to map protein interaction network *in situ* — isPIN.

## INTRODUCTION

Protein interaction network (PIN) or interactome has been mapped vigorously for the entire genome of *Drosophila* (Giot et al., 2003; Guruharsha et al., 2011) and human (Hein et al., 2015; Rolland et al., 2014). We recognize, nonetheless, that such a map could illuminate profound insights had its context been revealed. Since 1994, GFP (green fluorescent protein) (Chalfie et al., 1994) and other genetically encoded fluorescent proteins (Tsien, 1998) have labeled numerous proteins. They are useful in exposing inherent distribution biases of proteins within cells of diverse animals (Lukyanov, 2011), as well as serving as an immunological tag for purification of additional proteins capable of co-complexing (Guruharsha et al., 2011; Hein et al., 2015). In addition to these applications, valuable information regarding the way proteins interact *in vivo* can be collected using labeled proteins. Proximity attained by a pair of proteins that are labeled with spectrally overlapping fluorophores induces Förster resonance energy transfer (FRET)*, i.e.,* transfer of energy at near field from a fluorescence donor to a fluorescence acceptor (Förster, 1948). Therefore, occurrence of FRET as described best through quantum physics (Lakowicz, 2006) serves as a visual proxy for when and where proteins of interest interact with one another (Clegg, 2010). The idea of proximity-based scrutiny of interacting proteins (Sharifai et al., 2014) is not new, as the use of two-domain protein GAL4 transcription factor of yeast (Fields and Song, 1989) has expanded successfully to non-yeast proteins (Giot et al., 2003; Rolland et al., 2014). In contrast to yeast two-hybrid assays that sensitively demonstrate proteins’ binding compatibility, we maintain the native environment of the proteins we assess although, similarly to these assays, for a pair at a time. Our optimization aims for both the quality design in labeling proteins and the tight control in expressing labeled proteins. By using our protein labeling method (see Table1), it will be possible to re-supply natural context back to the map of protein interactome.

**Table 1.** Tools for charting freely interacting proteins within live animals.^‡^.

| **Tools** | **How** | **Why** | **Note** |
| --- | --- | --- | --- |
| (a) Genetically-encoded fluorescent proteins. | Fuse protein of interest (POI) to eGFP, mCherry or NirFP. | Visualize co-localization of proteins, and detect possible interaction between them by FRET. | Structural features of the β-barrel “cage” and flexible “loop” make eGFP, mCherry and NirFP reliable labels. |
| (b) Position-specific integration of transgene by PhiC31 integrase. | *attP* sequences allows integration of eGFP-labeled constructs to the *attP40* site and mCherry-labeled constructs to the *VK01* site on chromosome 2, and the NirFP-labeled constructs to the *attP2* site on chromosome 3. | Construct single, double and triple transgenic stocks through standard genetic crosses. | Landing sites selected are likely to be least problematic in terms of expressing labeled proteins. |
| (c) Expression control by GAL4 transcription factor. | Tandem *UAS* sequence allows for expressing transgenes under the control of a *GAL4* driver. | Control the time, place and dosage of expressing labeled proteins. | GAL4-responsive expressions typically cause no anatomical or behavioral anomalies. |

<sup>‡</sup> We integrate: (a) genetically encoded fluorescent proteins, (b) PhiC31 mediated genomic integration and (c) GAL4 responsive expression control. Use of these tools is not technically limited to *Drosophila*.

**Table 2.**
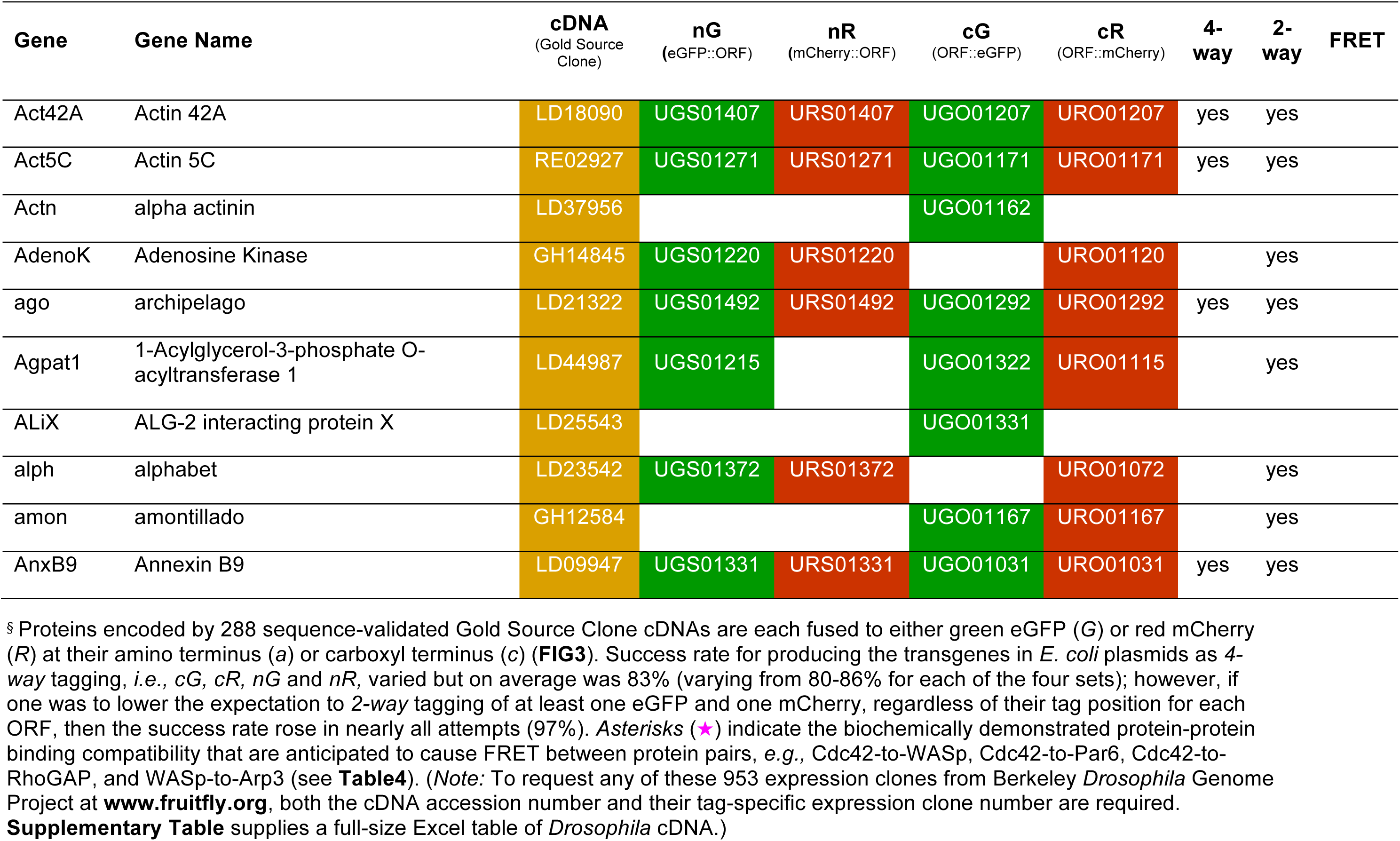
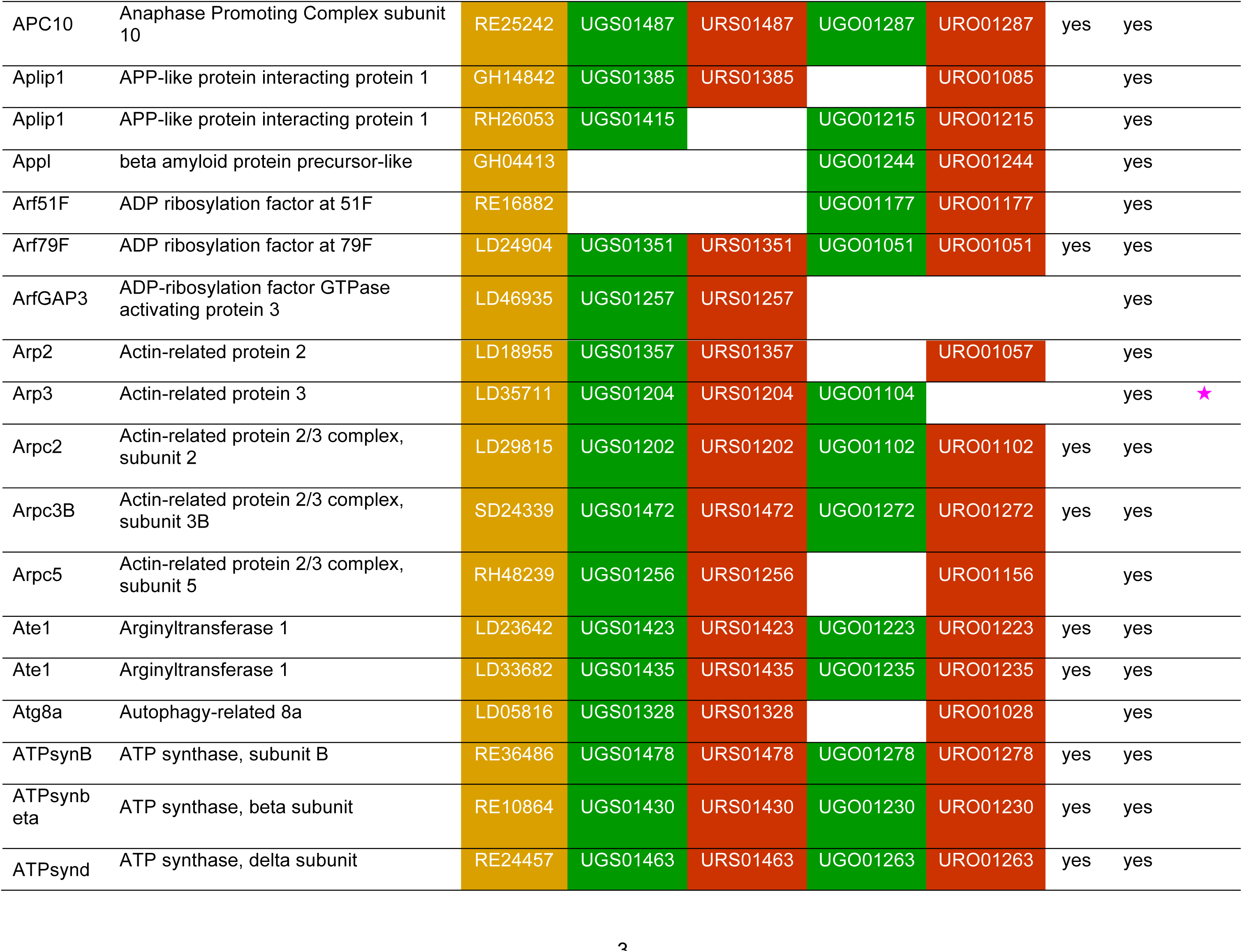
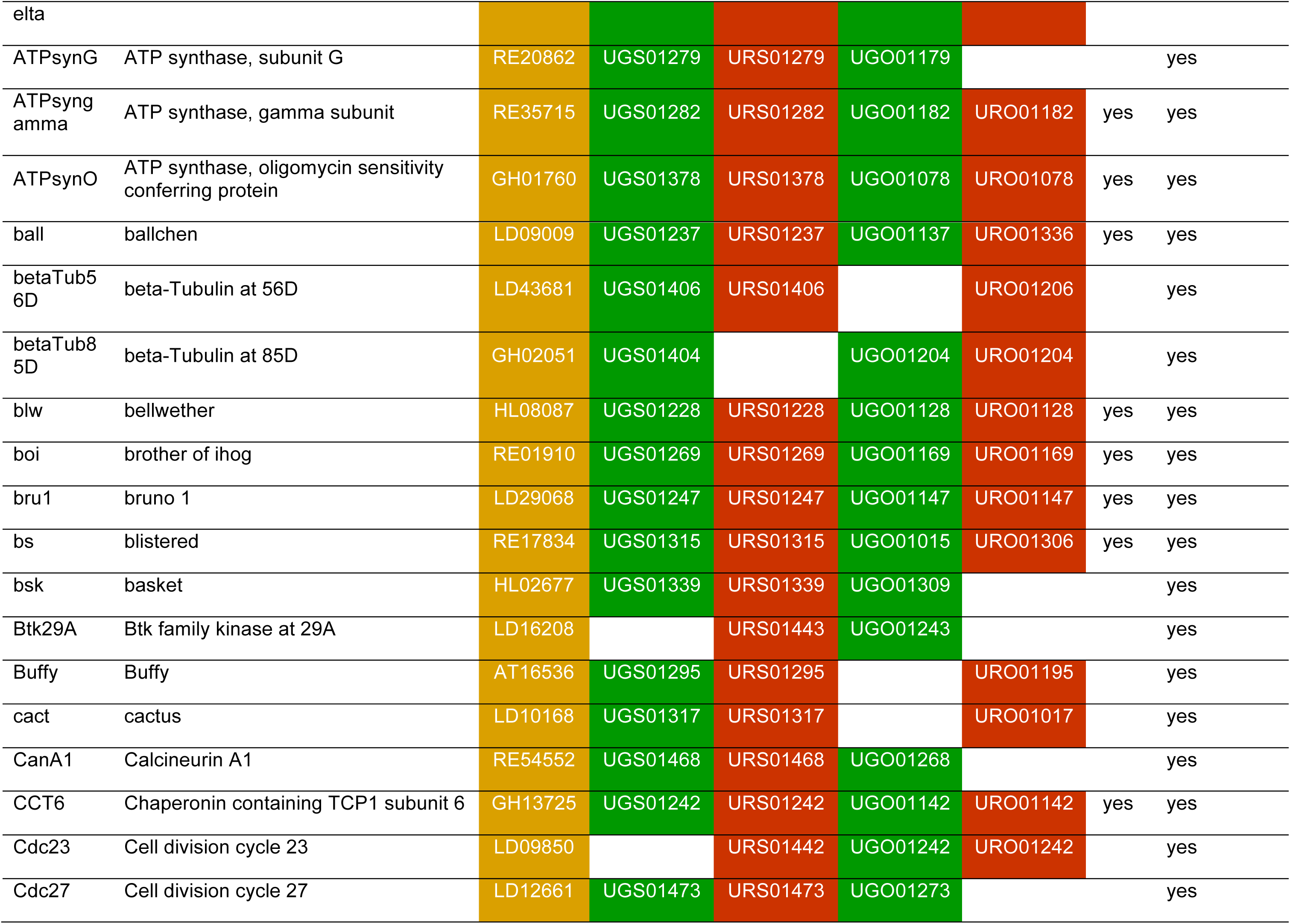
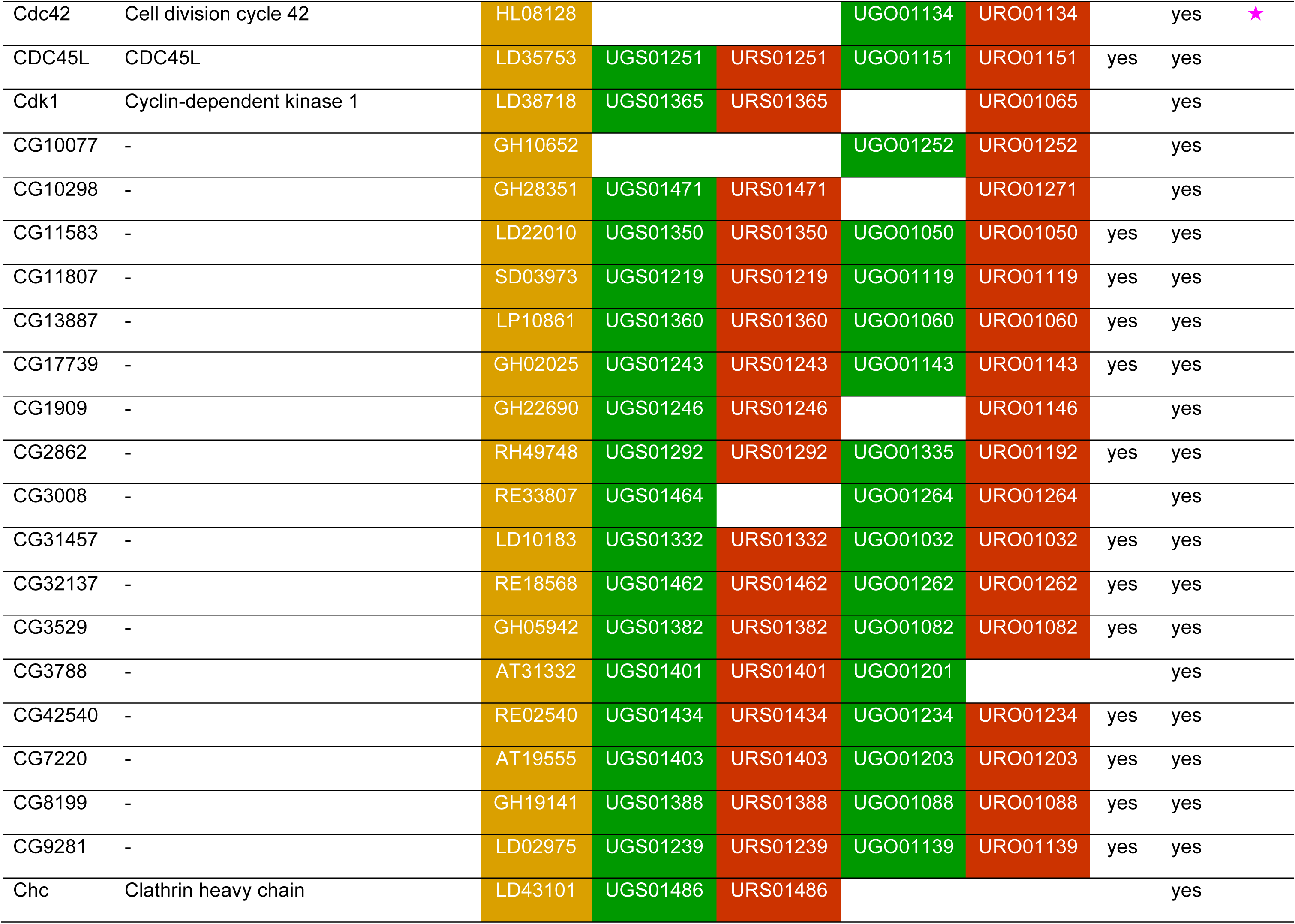
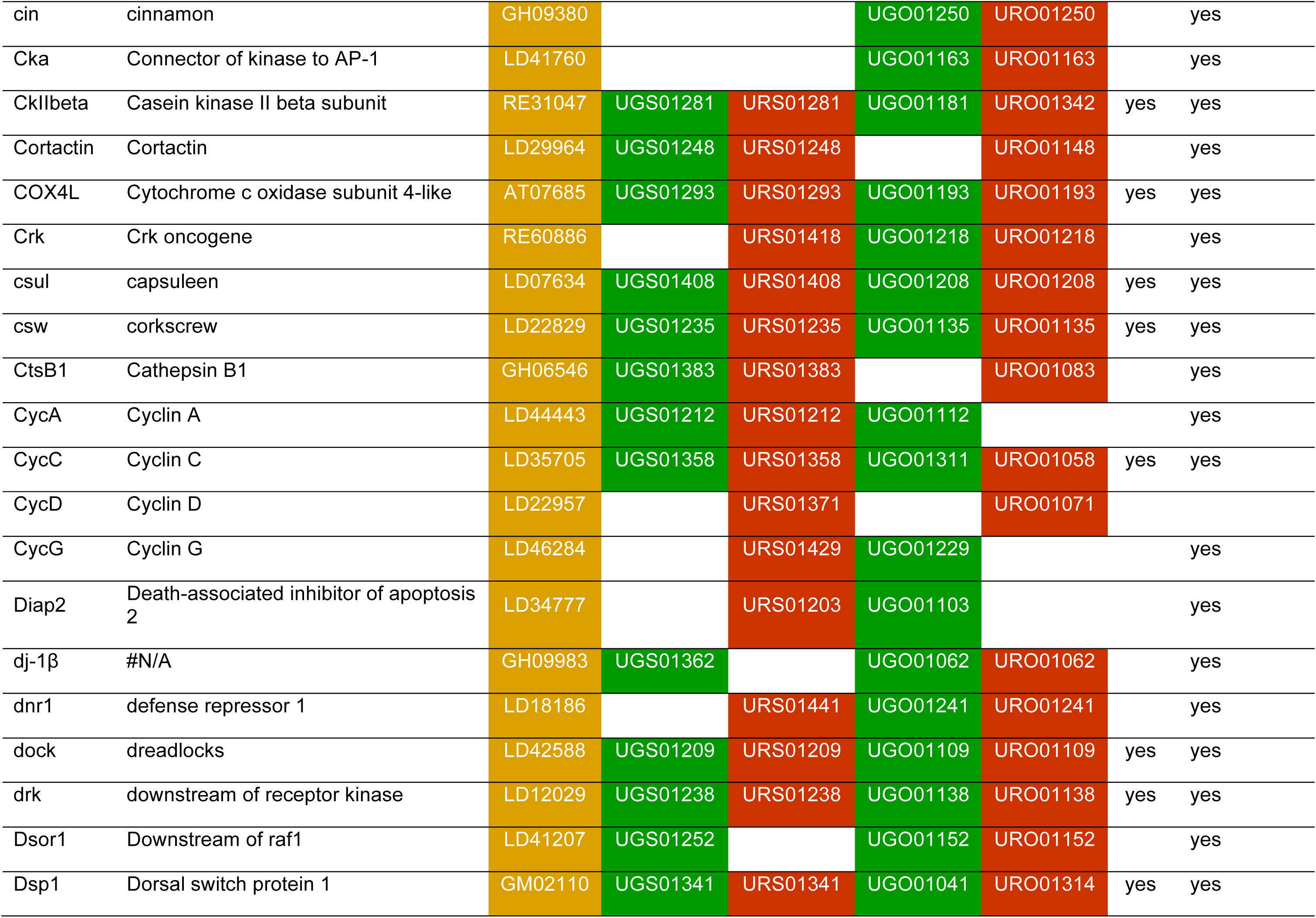
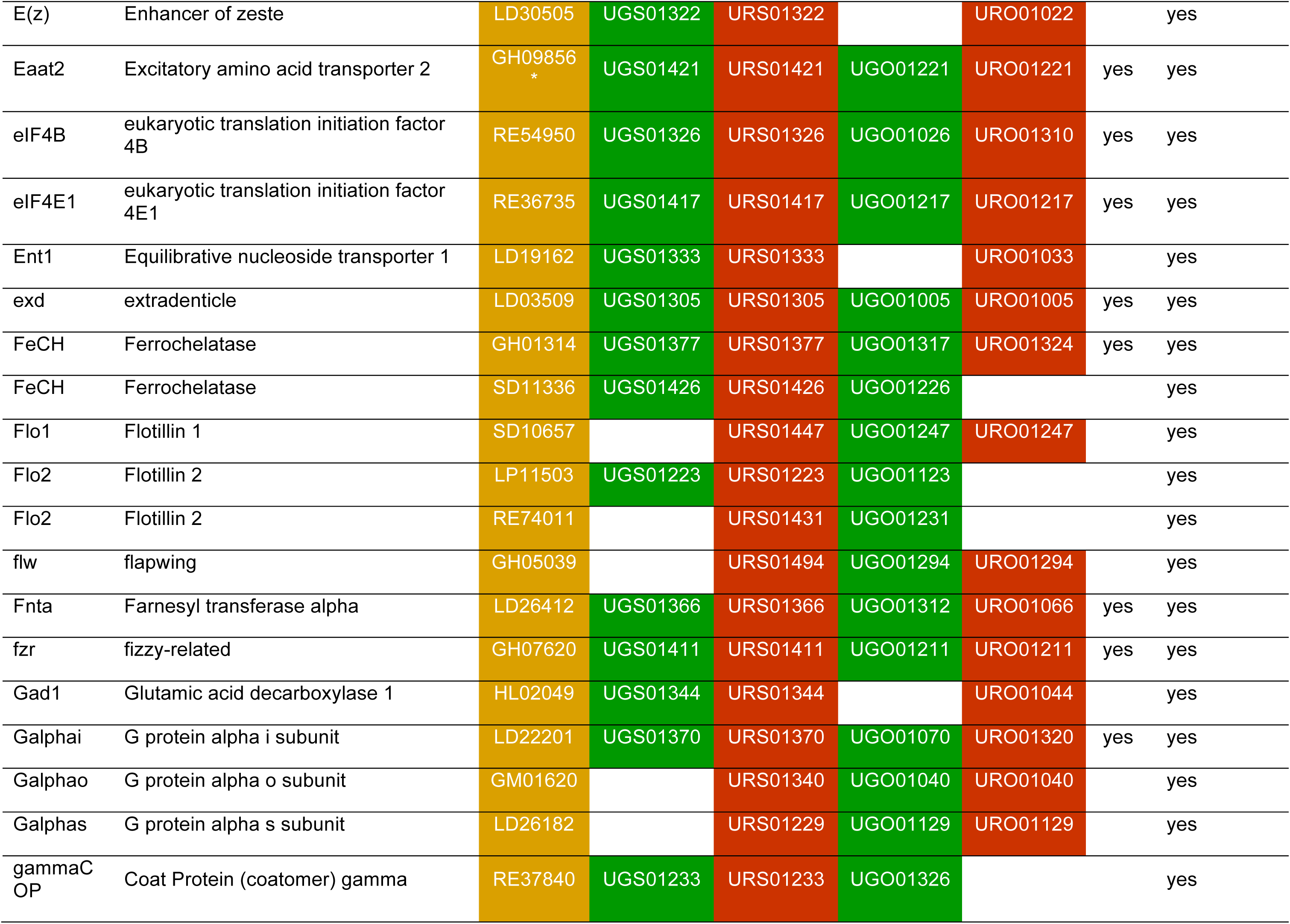
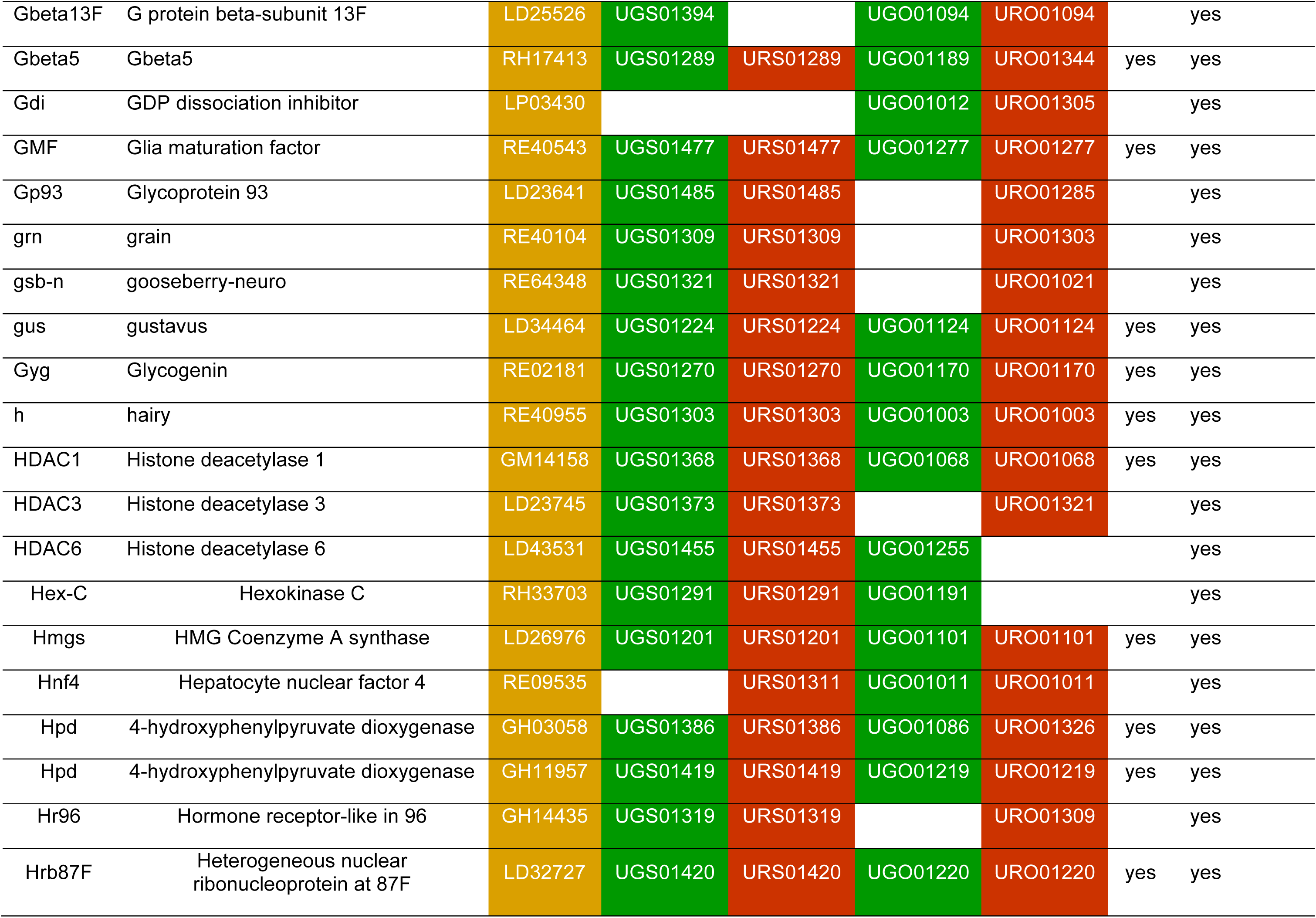
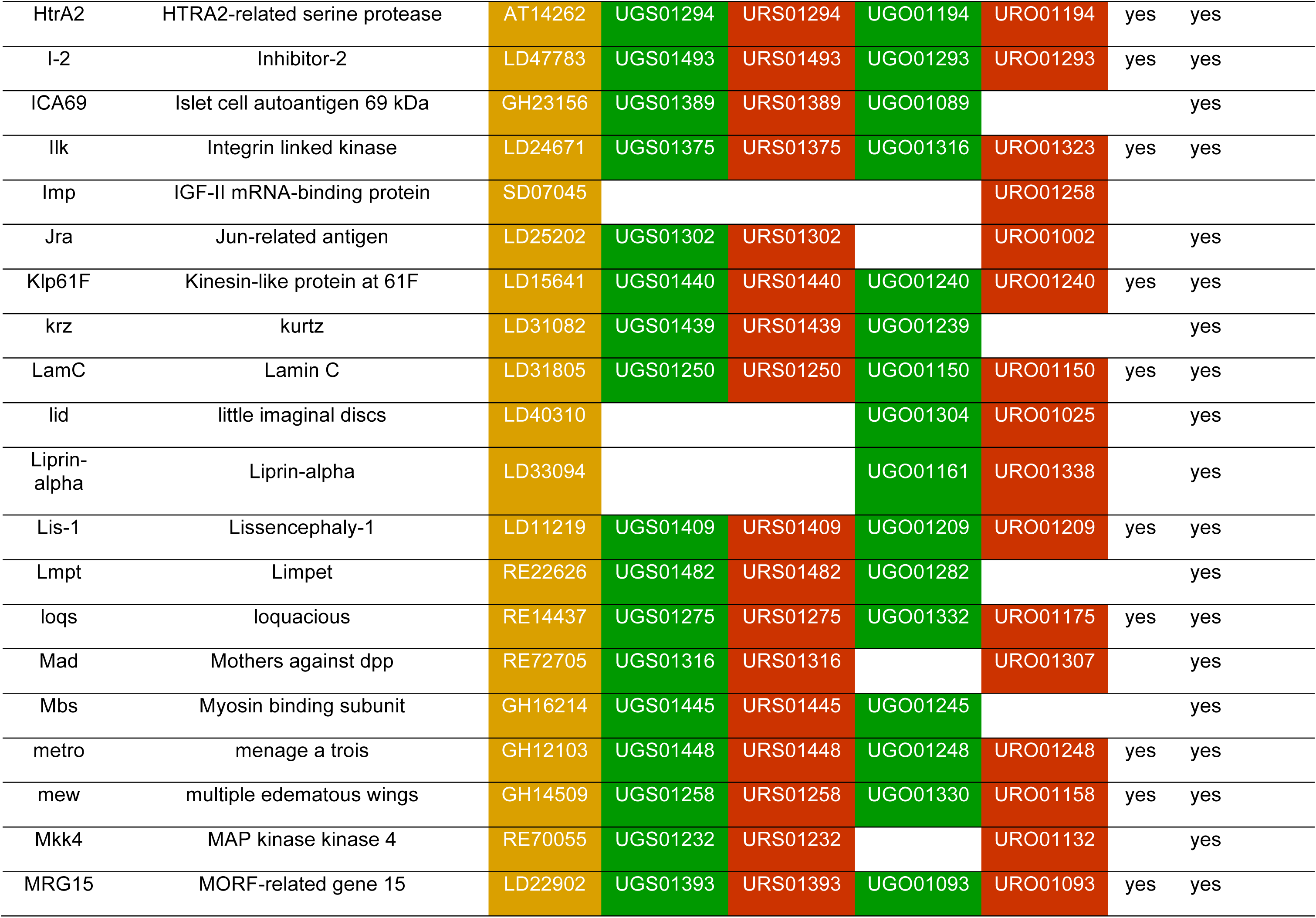
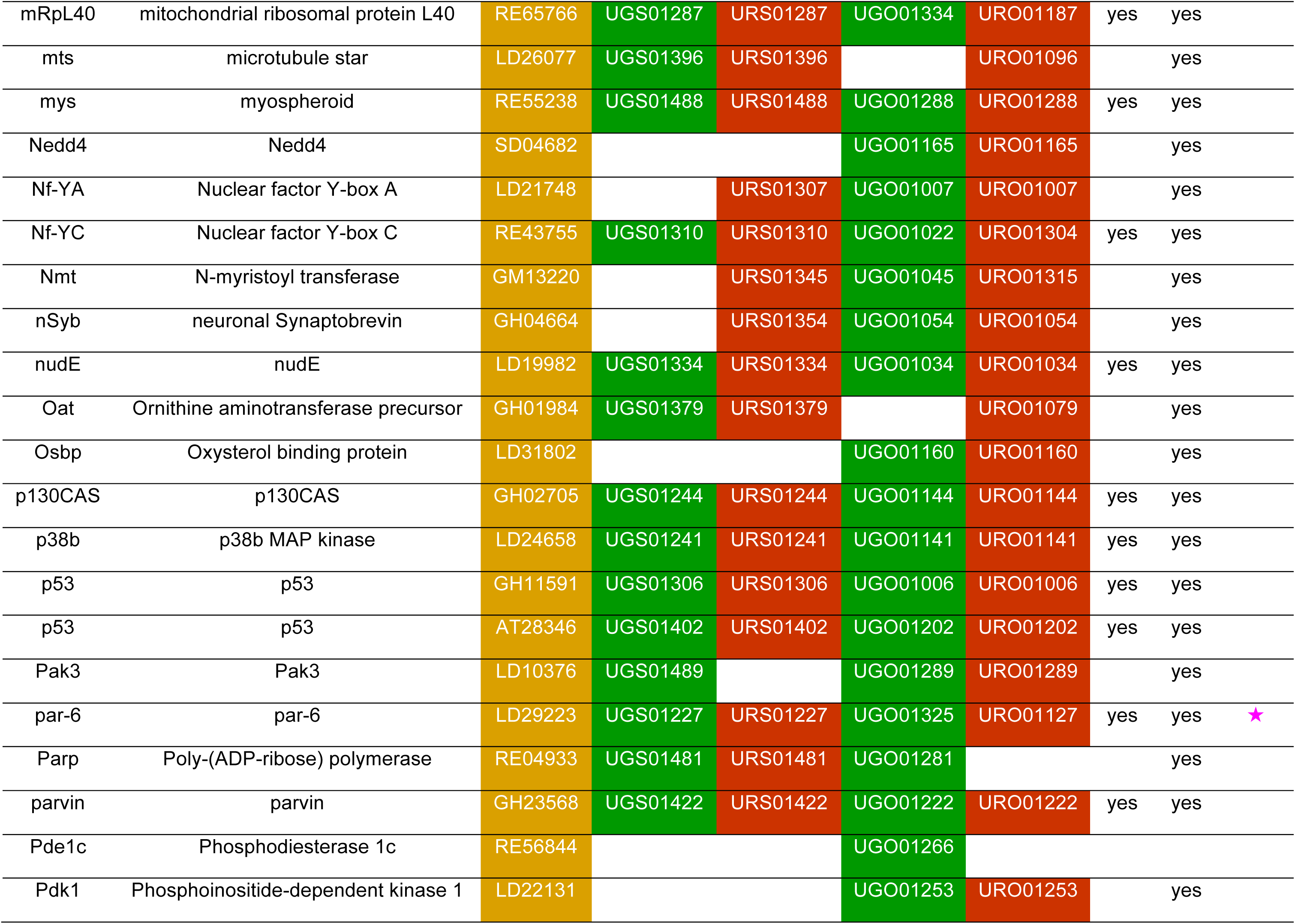
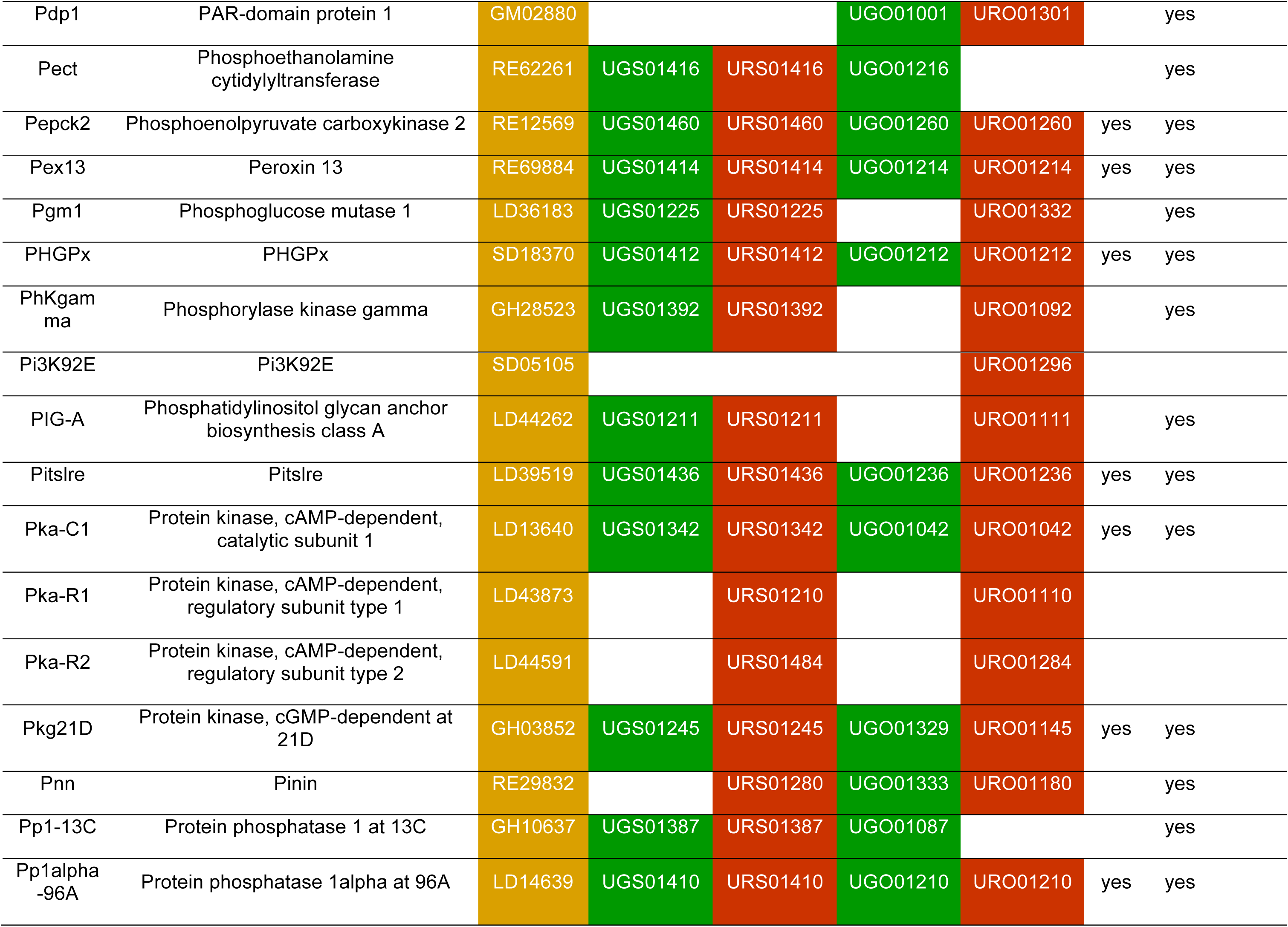
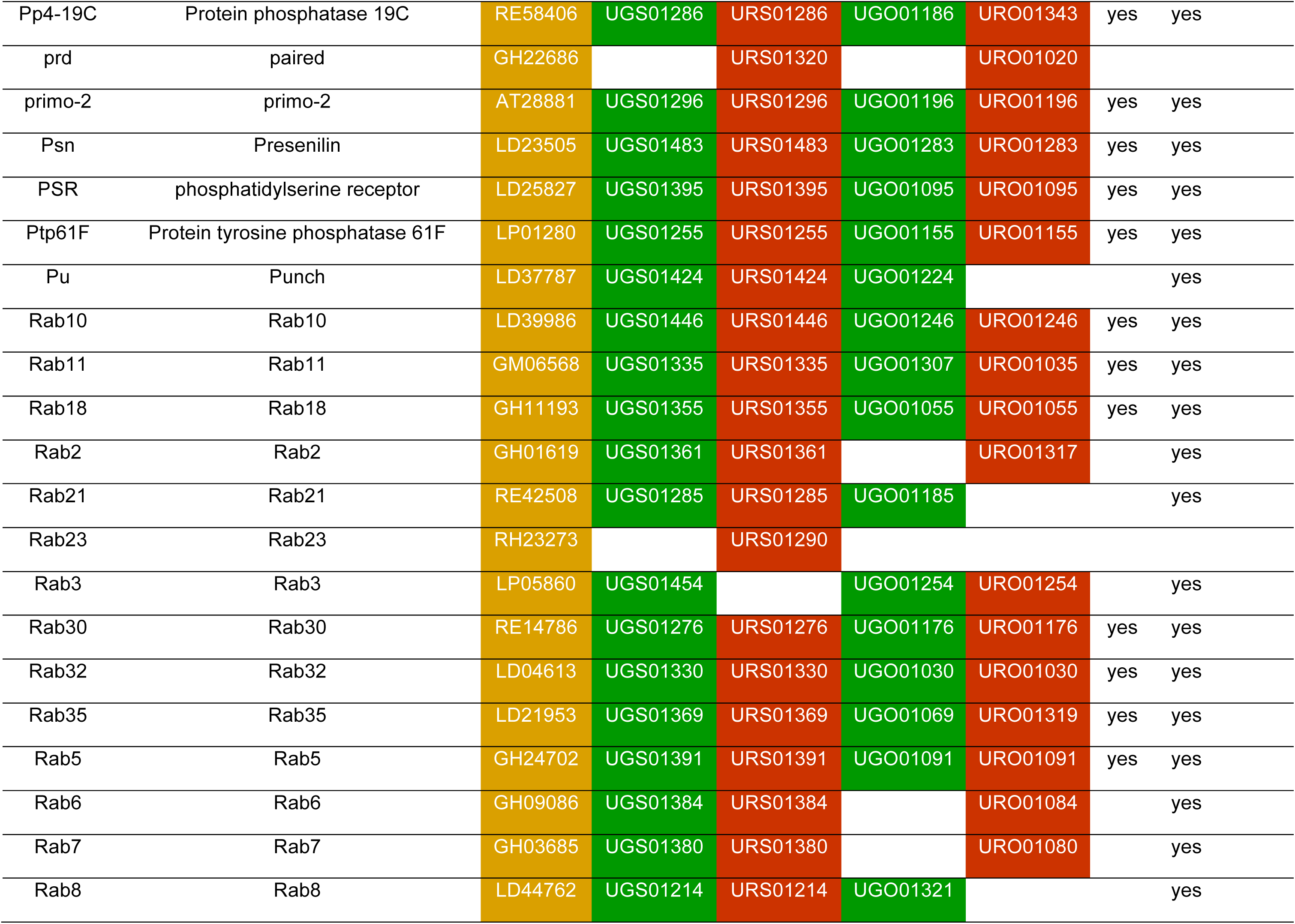
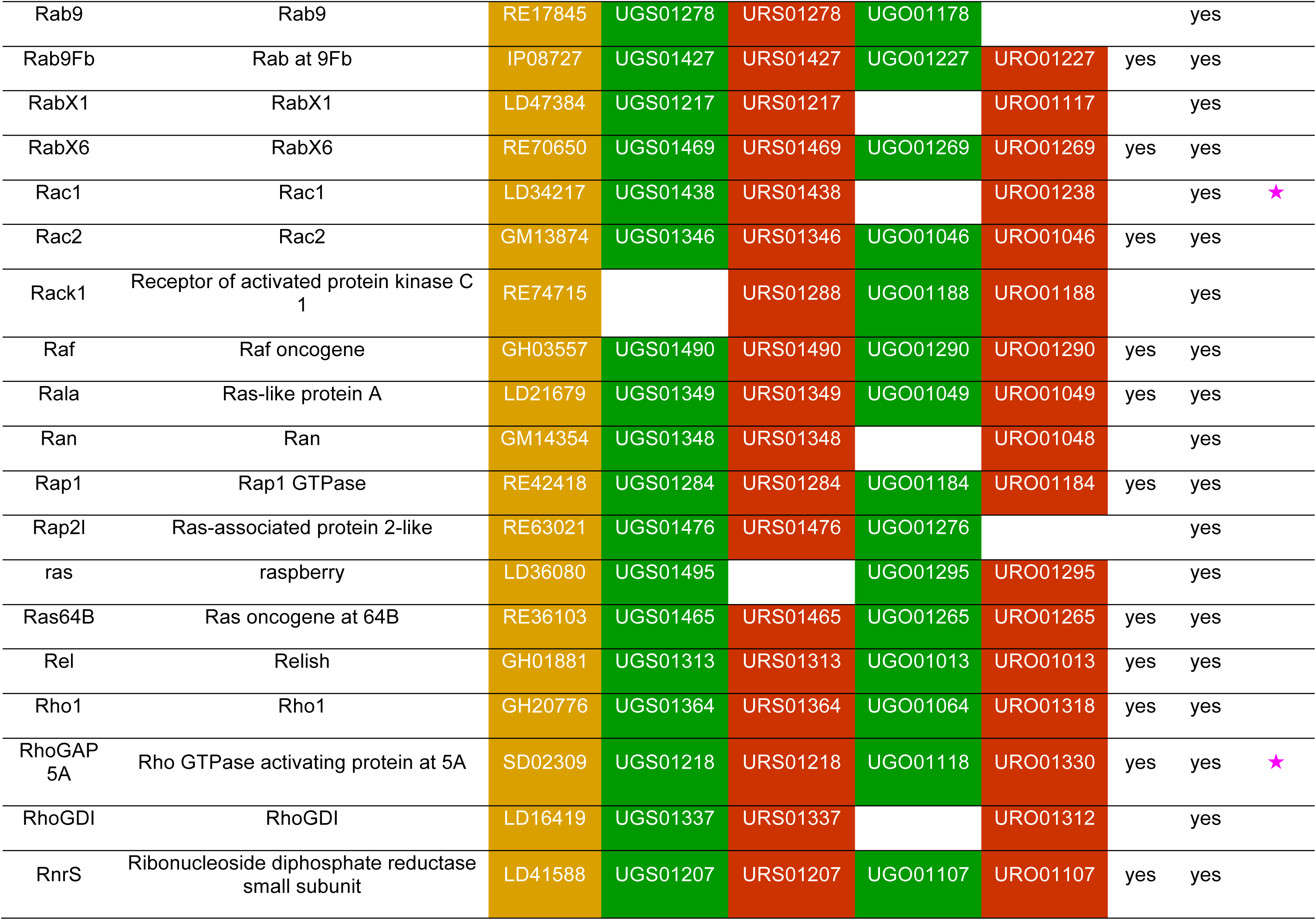
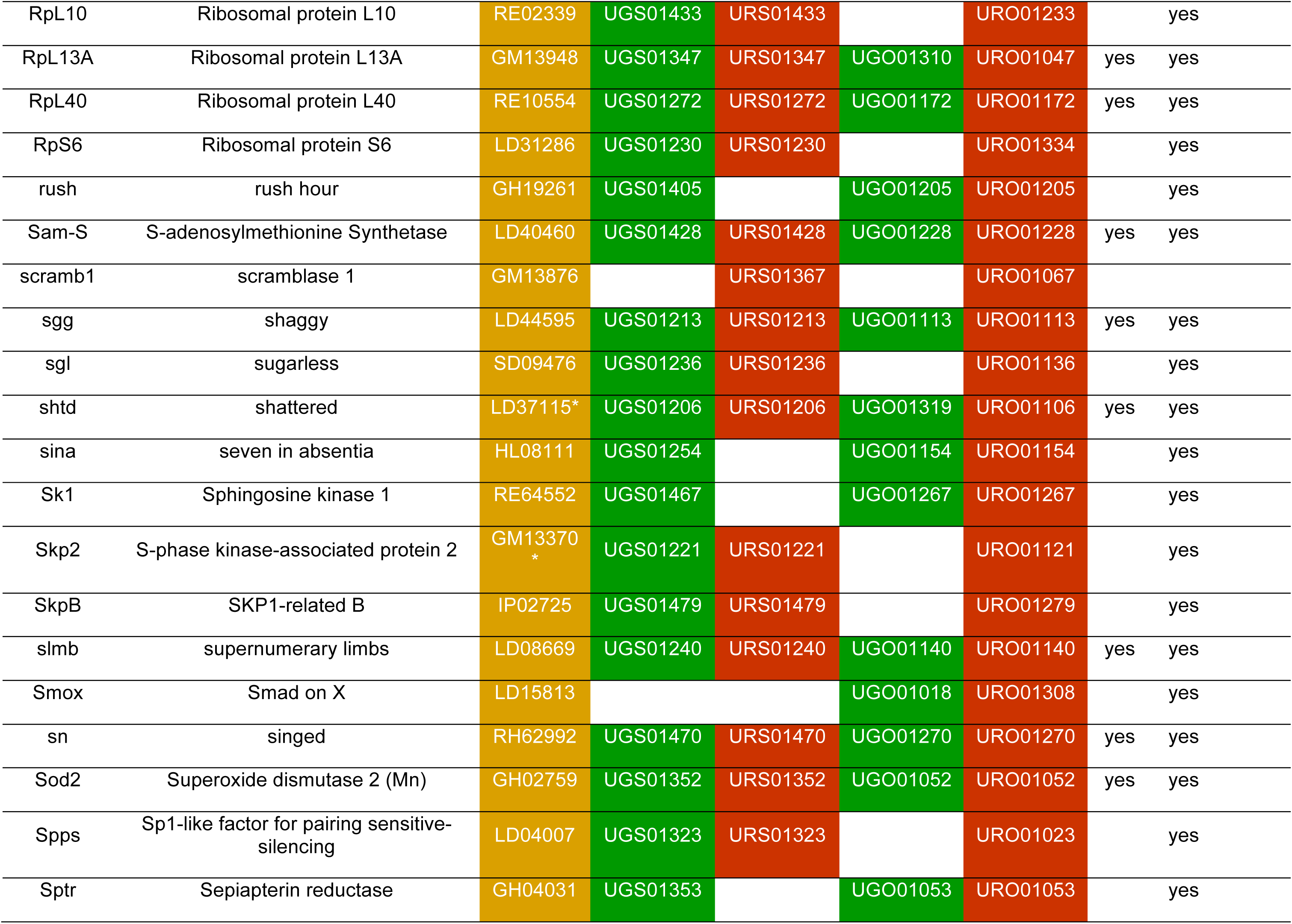
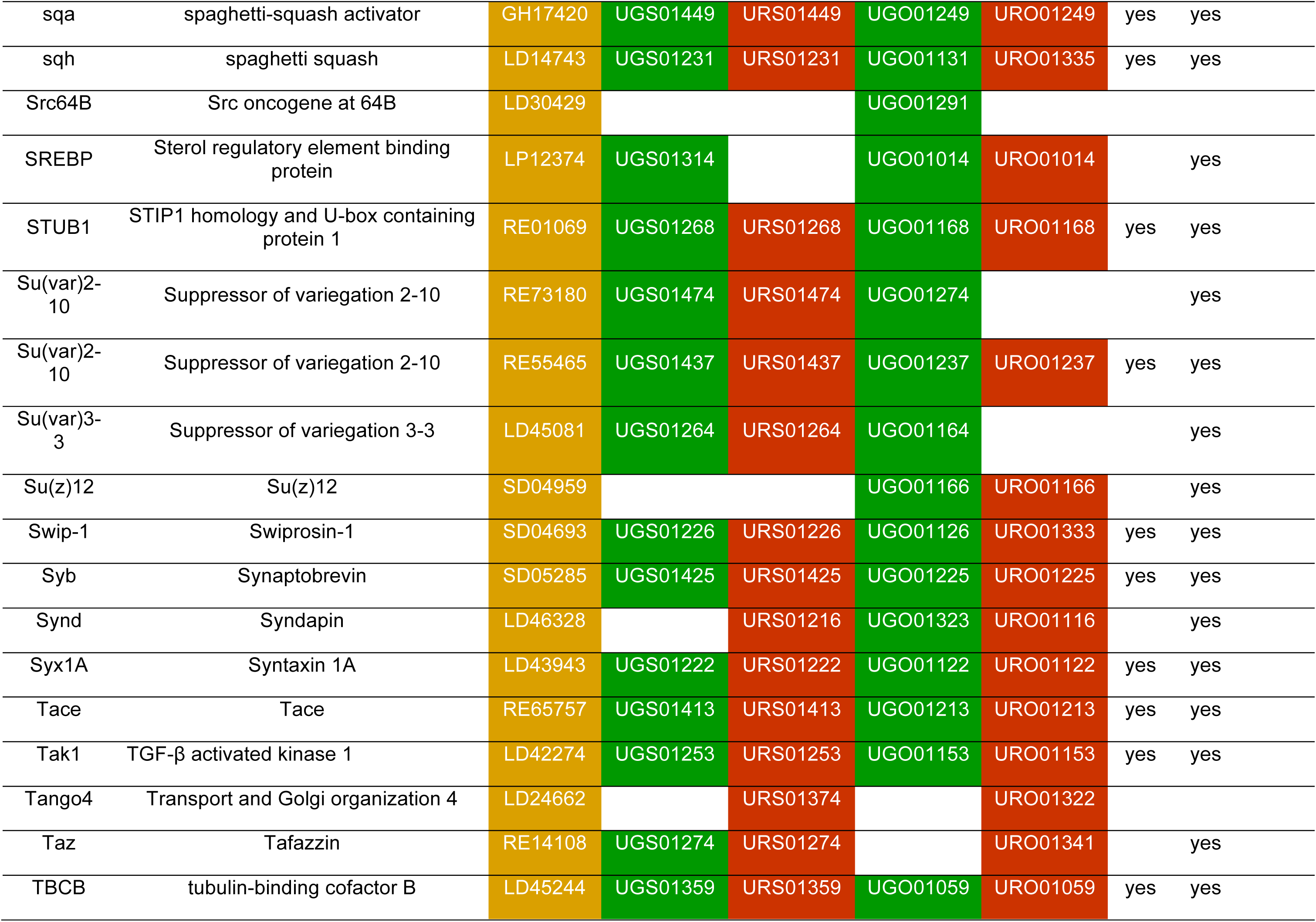
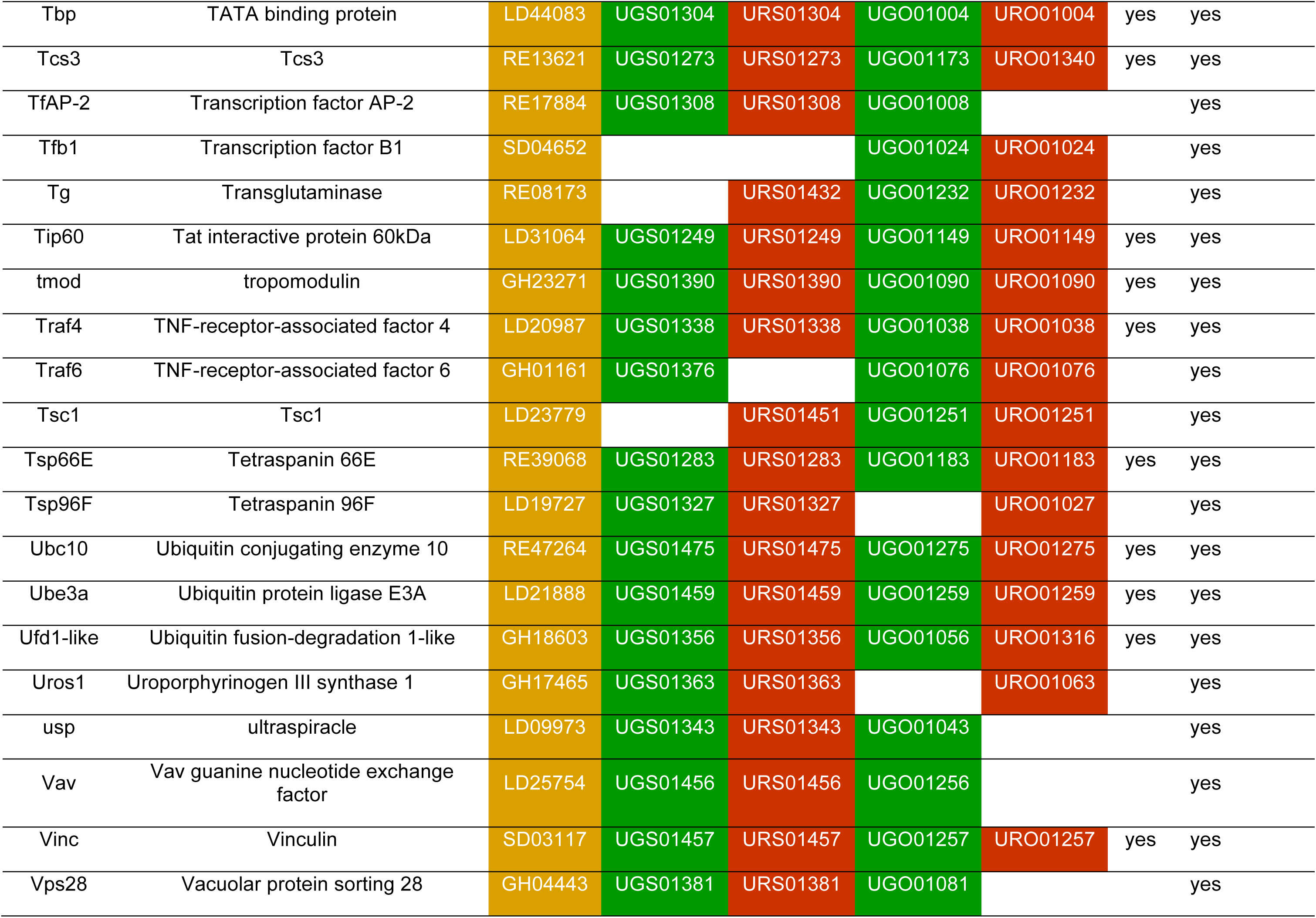
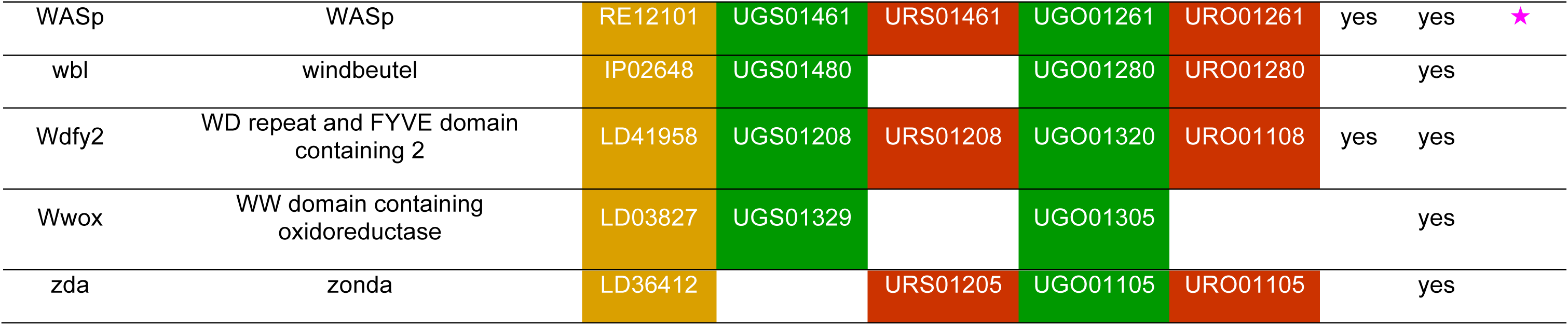
Drosophila cDNA. ^§^.

**Table 3.** Production steps.^**^

| Production steps | Attempted | Obtained | Success | Note |
| --- | --- | --- | --- | --- |
| (1) Construct production. | 286 proteins with 4-way labeling equals a total of 1144 expression constructs. | 988 expression constructs. | 83% for 4-way.<br>97% for 2-way. | 2-way labeling is most likely sufficient for checking possible interactions *in vivo*. |
| (2) Genome transformation. | 32 proteins (128 4-way labeled transgenes) selected from (1). | All 128 homozygous single transgenic stock lines. | 100%. | Transformation of transgenes into the *Drosophila* genome is straightforward. |
| (3) Stock construction. | Only 11 protein-protein links in (2) had their binding compatibility demonstrated biochemically; besides the Cdc42-to-WASp link (see Sharifai et al., 2014), the remaining 10 links were yet to be characterized *in vivo*. | 18 double homozygous and 4 triple homozygous stock lines (see **Table 4**). | 100%. | Establishing GAL4-responsive double and triple transgenic stock lines is straightforward. |
| (4) Sample visualization. | Triple transgenic stocks from (3) were crossed to a pan-neuronal *elav-GAL4* driver. Eggs were then collected and raised subsequently through larval stages. | Embryos developed normally without any apparent anomaly in anatomy or behavior until third instar larval stage. | 76%. | Nearly the same viability as in the previous report on double transgene stocks (Sharifai et al. 2014). |

\*\* In terms of production difficulty, transgene creation step (1) is the toughest and subsequent steps (2) through (4) are all forthright. In terms of time and cost of ultimately charting the map, however, step (4) can be the most extensive. Pre-selection of protein-protein links that are likely to produce FRET in the native environment is recommended.

**Table 4.**
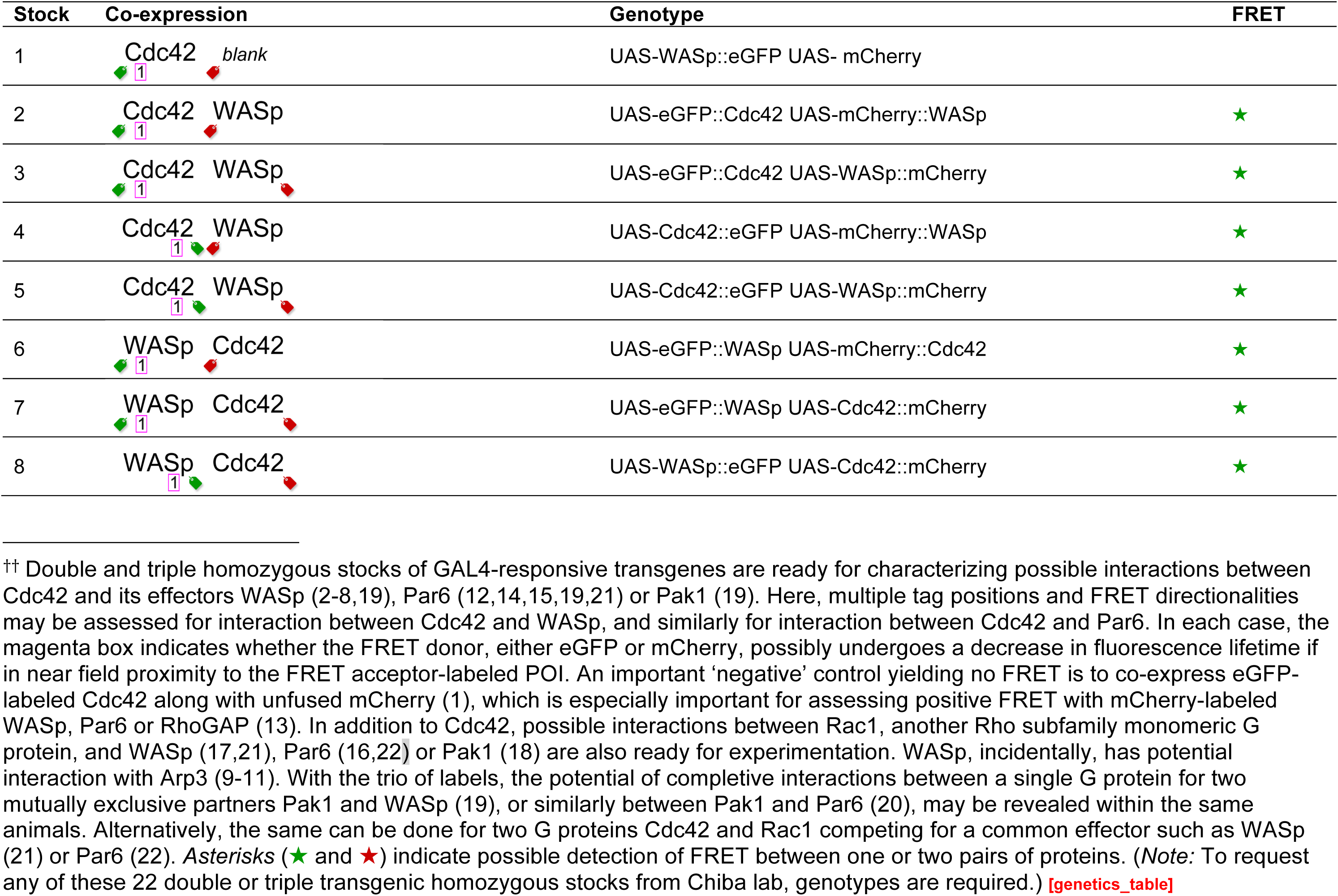
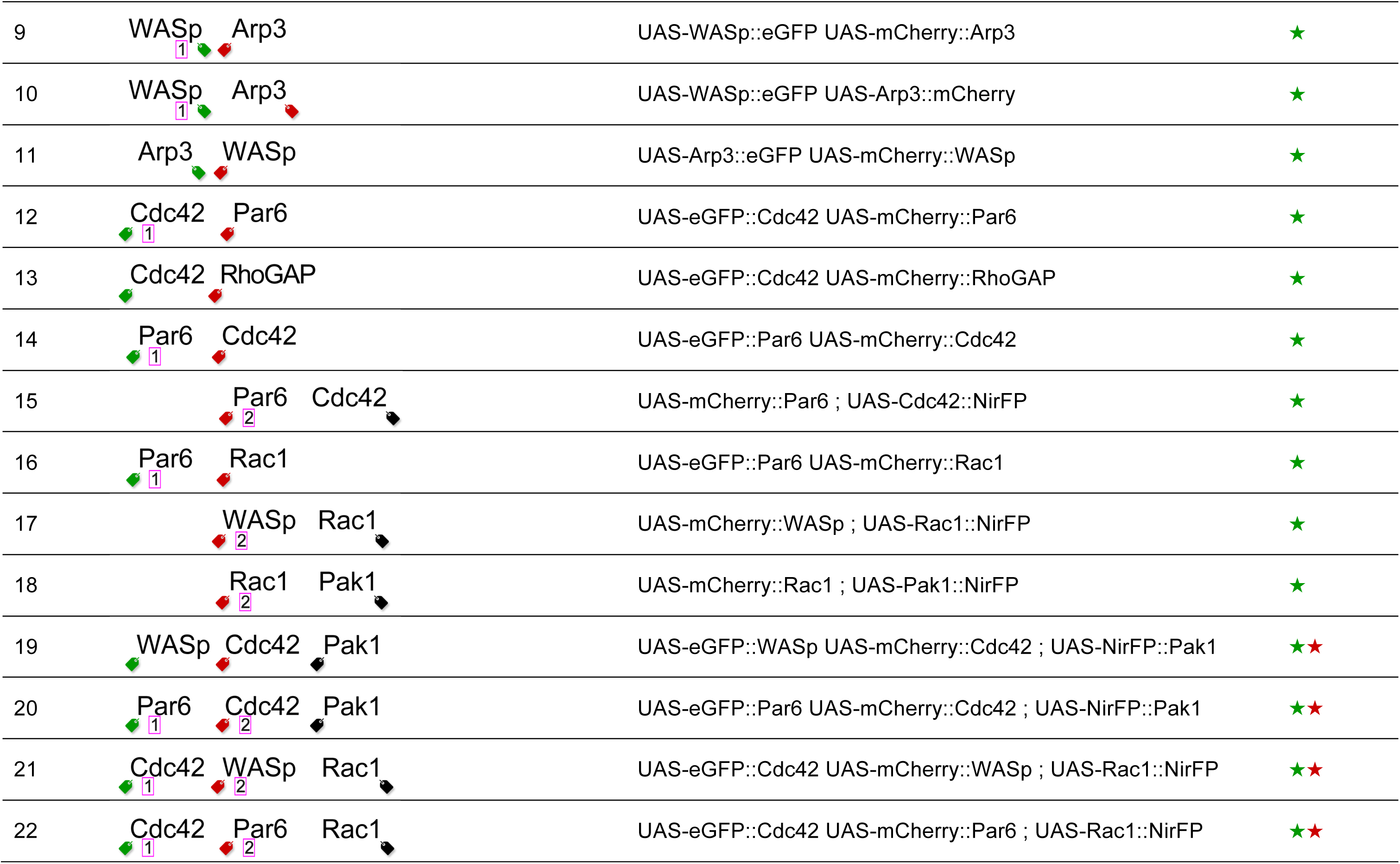
Drosophila stocks. ^††^.

| Stock | Co-expression | Genotype | FRET |
| --- | --- | --- | --- |
| 1 | Cdc42 blank | UAS-WASp::eGFP UAS- mCherry |  |
| 2 | Cdc42 WASp | UAS-eGFP::Cdc42 UAS-mCherry::WASp | ★ |
| 3 | Cdc42 WASp | UAS-eGFP::Cdc42 UAS-WASp::mCherry | ★ |
| 4 | Cdc42 WASp | UAS-Cdc42::eGFP UAS-mCherry::WASp | ★ |
| 5 | Cdc42 WASp | UAS-Cdc42::eGFP UAS-WASp::mCherry | ★ |
| 6 | WASp Cdc42 | UAS-eGFP::WASp UAS-mCherry::Cdc42 | ★ |
| 7 | WASp Cdc42 | UAS-eGFP::WASp UAS-Cdc42::mCherry | ★ |
| 8 | WASp Cdc42 | UAS-WASp::eGFP UAS-Cdc42::mCherry | ★ |
| 9 | WASp Arp3 | UAS-WASp::eGFP UAS-mCherry::Arp3 | ★ |
| 10 | WASp Arp3 | UAS-WASp::eGFP UAS-Arp3::mCherry | ★ |
| 11 | Arp3 WASp | UAS-Arp3::eGFP UAS-mCherry::WASp | ★ |
| 12 | Cdc42 Par6 | UAS-eGFP::Cdc42 UAS-mCherry::Par6 | ★ |
| 13 | Cdc42 RhoGAP | UAS-eGFP::Cdc42 UAS-mCherry::RhoGAP | ★ |
| 14 | Par6 Cdc42 | UAS-eGFP::Par6 UAS-mCherry::Cdc42 | ★ |
| 15 | Par6 Cdc42 | UAS-mCherry::Par6 ; UAS-Cdc42::NirFP | ★ |
| 16 | Par6 Rac1 | UAS-eGFP::Par6 UAS-mCherry::Rac1 | ★ |
| 17 | WASp Rac1 | UAS-mCherry::WASp ; UAS-Rac1::NirFP | ★ |
| 18 | Rac1 Pak1 | UAS-mCherry::Rac1 ; UAS-Pak1::NirFP | ★ |
| 19 | WASp Cdc42 Pak1 | UAS-eGFP::WASp UAS-mCherry::Cdc42 ; UAS-NirFP::Pak1 | ★ ★ |
| 20 | Par6 Cdc42 Pak1 | UAS-eGFP::Par6 UAS-mCherry::Cdc42 ; UAS-NirFP::Pak1 | ★ ★ |
| 21 | Cdc42 WASp Rac1 | UAS-eGFP::Cdc42 UAS-mCherry::WASp ; UAS-Rac1::NirFP | ★ ★ |
| 22 | Cdc42 Par6 Rac1 | UAS-eGFP::Cdc42 UAS-mCherry::Par6 ; UAS-Rac1::NirFP | ★ ★ |

## RESULTS

To label individual proteins, we chose genetically encoded fluorescent proteins eGFP (Bierhuizen et al., 1997), mCherry (Shaner et al., 2004) and NirFP (Shcherbo et al., 2010) (FIG1). These labels serve as versatile imaging tools to study proteins within live animals. First, each of these fluorophores has structural resemblance to a “cage”. This cage-like configuration is likely the strongest natural means for protecting a set of fluorescing atoms from surrounding stress. More specifically, the extremely rare ß sheet barrels constituting these proteins are proposed to solidly safeguard a small α helix, T65-Y66-G67 in eGFP for instance, capable of exhibiting fluorescent properties (Ormo et al., 1996). Another well-known example of ß sheet barrels protecting delicate internal structures occurs among channel proteins, such as in a tetrameric potassium channel (Zhou et al., 2001), that regulate ion selectivity of the cell surface. Second, each of these fluorescent labels is physically linked to a protein of interest through a “loop.” Unlike synthetic chemicals that need conjugation through much weaker non-covalent bonds, this robustly covalent yet non-structural linkage achieves two consequences: (*1*) a short and constant distance between the label and the protein that is labeled, and (*2*) randomization of dipole orientation known as *κ****^2^*** of the label with respect to the protein it labels. Unlike synthetic fluorescent dyes (Pertz et al., 2006) or fluorescence-absorbing metal atoms (Koch and Larsson, 2005), neither eGFP, mCherry, nor NirFP requires pre-imaging dissection or incubation of animals after attaining full maturation, typically within 2-4 hours. As mentioned elsewhere (Chong et al., 2015; Hudson et al., 2008; Sarov et al., 2012; Trinh le et al., 2011; Venken et al., 2011), these and other genetically encoded fluorescent proteins are extremely valuable for time lapse imaging of proteins during development and behavior of animals. eGFP, mCherry and NirFP are derived from jellyfish *Aequoria* and sea anemone, *Discosoma* and *Entacmaea,* respectively. Unlike their natural forms, however, custom-tuned eGFP and mCherry are each monomeric and do not dimerize by themselves. Also important, these two and NirFP do not bind one another directly when co-expressed in model organisms such as *Drosophila* (our observation). This is satisfactory because during imaging, both eGFP and mCherry are required to serve as a FRET donor. Any inherent affinity, whether previously known or unknown, between labeled proteins needs to be the sole reason for inducing FRET. Absorption and emission spectra of these fluorescent labels are distinct from each other. Notably, the eGFP-mCherry-NirFP trio offers two excellent yet separable FRET pairs: eGFP-to-mCherry with a Förster distance of 5.2 nm and mCherry-to-NirFP with a Förster distance of 4.4 nm. Multiplexing FRET capabilities using this trio may therefore serve as the basis for exposing the logic governing and/or the extent delimiting proteins interacting with each other (not shown). Although we are not aware of any systematic test on the functional neutrality of protein labeling, we believe that genetic knockouts of any specific proteins, if attempted, may be rescued by using fluorescently labeled but otherwise wild type forms of proteins. In fact, the very first use of GFP in fruit flies reported that genetic knockout of a certain nuclear protein could be rescued by expressing a GFP-fused wild-type form (Wang and Hazelrigg, 1994). More recently, Cdc42 protein, required for initiating the dendrites in neurons (Kamiyama and Chiba, 2009), was shown to rescue animals lacking a *cdc42* gene, regardless of whether the wild type transgene was fused to eGFP or not (our observation). We did not adopt split GFP as in the GRASP method (Feinberg et al., 2008) or sequential two-step GFP formation as in the GFP11 knock-in method (Kamiyama et al., 2016) because these strategies do not distinguish transient from permanent interactions. It should be further noted that these fluorescent proteins cannot readily fuse to DNA or RNA covalently, thus limiting their potential usage for exposing binding specificity of certain transcription factors or other DNA-binding/RNA-binding proteins toward particular nucleotide sequences through FRET.

**Figure 1.**
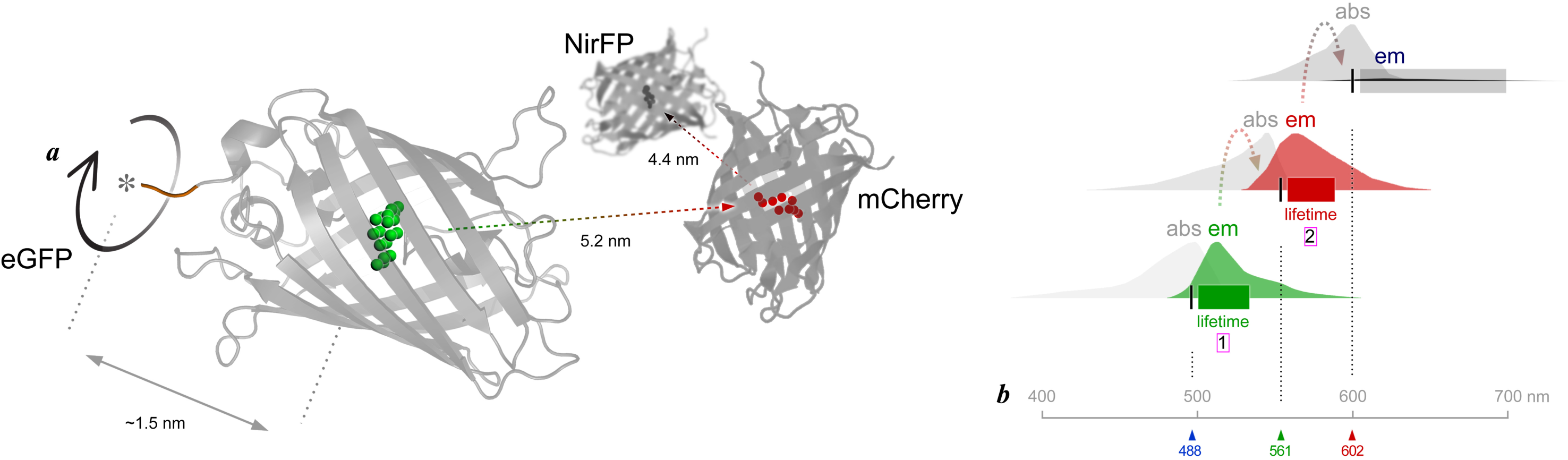
Labels. (***a***) Structures of fluorescent labels. Monomeric eGFP has twenty-two atoms (green) fluoresce together at the center of a “cage” — a rigid barrel structure composed of eleven anti-parallel ß sheets.The distance from the center of this α helical fluorophore to the edge of a protein of interest (gray *asterisk*) is about 1.5 nm. eGFP’s carboxyl terminus with an additional 17-mer polypeptide (brown) exhibits a great deal of flexibility, (circle *arc*). This “loop” is anticipated to randomize the polar orientation *κ^2^* of eGFP in respect to the protein it labels, thereby dramatically simplifying the computation of Förster resonance energy transfer that may occur between different fluorescent labels. The emission of eGFP and the absorption of mCherry display extensive spectral overlap (see *(b)*), providing a possibility of FRET with the Förster distance of 5.2 nm between these two fluorophores. Monomeric mCherry also has fluorescing ions (red) surrounded by a barrel composed of anti-parallel ß sheets. Its emission and the absorption of NirFP display generous spectral overlap (see *(b)*) and provide a possibility of FRET with a Förster distance of 4.4 nm between these two fluorophores. The detailed structure of NirFP remains currently unknown. (***b***) Spectra of fluorescent labels. With a 488 nm diode laser (blue *arrowhead*), eGFP fluorescence (green *bar*) may be examined with its lifetime *τ* of 2.56 ns. Its emission (green *spectrum*) and the absorption of mCherry (gray *spectrum*) overlap. If any physical interaction between proteins *A* and *B* occurs, this induces FRET between the two fluorophores. Hence, shortening of the lifetime *τ* of the FRET donor eGFP (magenta *box1*) signifies FRET. With a 561 nm diode laser (green *arrowhead*), mCherry fluorescence (red *bar*) may be examined with its normal lifetime *τ* of 1.47 ns. Its emission (red *spectrum*) and the absorption of NirFP (gray *spectrum*) overlap. If any physical interaction occurs between proteins *B* and *C,* this induces FRET between the two fluorophores. With a 602 nm diode laser (red *arrowhead*), NirFP fluorescence (gray *bar*) may be examined. The crystal structure of NirFP is currently unavailable.

We developed expression vectors, pUAS-C-eGFP-BD-attB, pUAS-C-mCherry-BD-attB, pUAS-N-eGFP-BD-attB and pUAS-N-mCherry-BD-attB for generating fluorescently tagged proteins of interest (POIs) (FIG2). First, we transferred protein open reading frames (ORFs) to the expression vectors using the Cre-Lox recombination cloning system, similar to the strategy used to make FLAG-HA constructs for tandem-affinity purification conducted previously (Guruharsha et al., 2011). The use of 100% sequence-validated XO and XS clone sets (Yu et al., 2011) prevents any unknown or non-intentional modification of the proteins being analysed. For 286 selected genes we used 283 BO clones to produce carboxyl terminal fusions from the XO clone set and 264 BS clones to produce amino terminal fusions from the XS clone set. We generated 953 expression clones in total: 236 UGS (eGFP amino tag), 245 URS (mCherry amino tag), 228 UGO (eGFP carboxyl tag) and 244 URO (mCherry carboxyl tag). Second, the *attp* sequence added to each expression vector is required for targeted integration of transgenes into specific landing sites in the *Drosophila* genome. We used phiC31 integrase, as originally characterized in retroviruses (Groth et al., 2004). The landing sites selected, *attp40* and *vk01* on chromosome 2 and *attp2* on chromosome 3, have high integration rates and are also known to induce sufficiently high rates of expression as GAL4-reponsive transgenes (Markstein et al., 2008; Venken et al., 2006a). It is also straightforward to introduce two transgenes onto one chromosome as a double transgene stock though intentional homologous recombination. Third, a tandem *UAS* sequence added to the vectors allows for expressing transgenes under the control of yeast GAL4 transcription factor (Brand and Perrimon, 1993). This enhancer hijacking in a foreign host genome*, i.e., Drosophila* in our case, offers total control over the time and place for co-expressing up to three transgenes all at a constant dosage.

**Figure 2.**
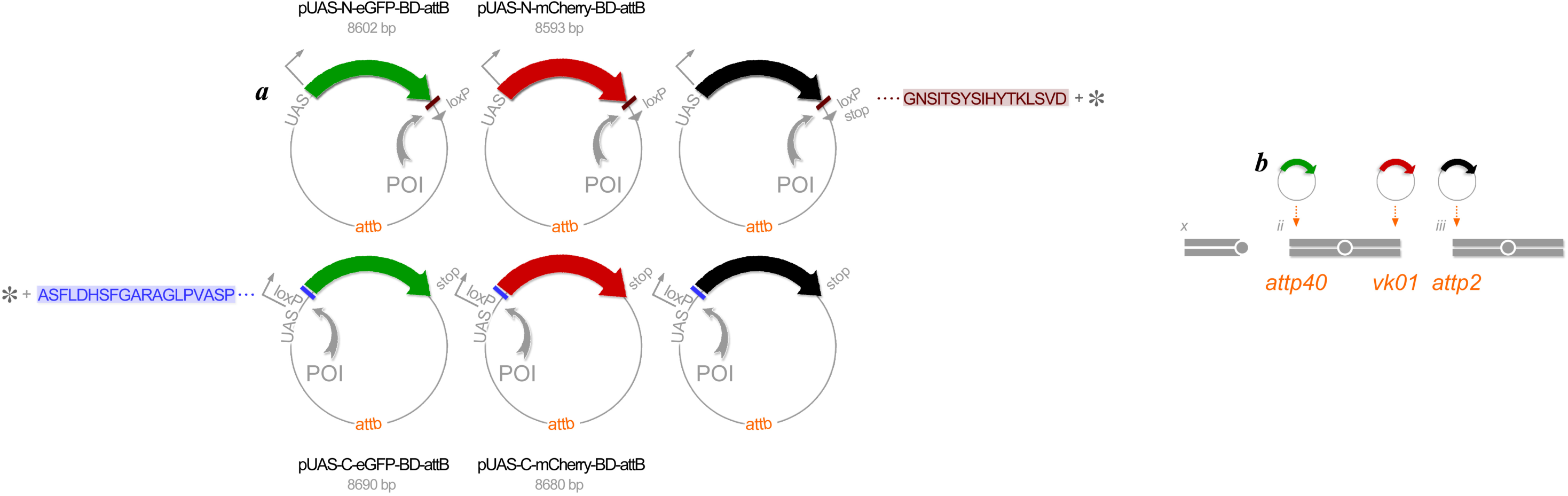
Transformation. (***a***) Expression vectors. Constructed in *E. coli* plasmids in six complementary manners, an individual protein of interest (POI) (gray *arc*) is labeled at either its amino terminus (top row) or its carboxyl terminus (bottom row) with eGFP (green), mCherry (red) or NirFP (black). A short, flexible polypeptide sequence (brown or purple *bar*) between the protein (gray *asterisk*) and its fluorescent label is added to randomize the dipole orientations when a pair of labeled proteins are being assessed for possible interaction via FRET occurrence. All these expression vectors are placed downstream of the *UAS* sequence for expressing the labeled proteins under the GAL4 expression system, and also carry the *attb* sequence for PhiC31-mediated genomic integration to specific landing sites (see *(b)*). All plasmids carry mini *w^+^* as a transformation marker (not shown). (***b***) Genomic integration. PhiC31 integrase enables position-specific genomic integration of all transgenes for eGFP-labeled proteins to the landing site *attp40* on the second chromosome, all transgenes for mCherry-labeled proteins to the landing site *vk01* on the chromosome 2, and all transgenes for NirFP-labeled proteins to landing site *attp2* on the chromosome 3.

Using this position-specific integration in the *Drosophila* genome, various single transgene, double transgene, and triple transgene stocks can be created (FIG3a-c). Homologous recombination occurs in the female germ line and, therefore, the use of balancer chromosomes during standard genetic crosses suppresses unwanted recombination of two transgenes in a single chromosome, *e.g.,* chromosome *ii*. As illustrated schematically, intentional uses of balancers afford stress-free combination of multiple transgenes, with all undergoing expression at constant dosage. Incidentally, a single genetic dosage is sufficient for detecting fluorescence from all labeled proteins, as well as detecting FRET quantitatively. The low-dosage expression also minimizes photo-toxicity during repeated and/or time-lapse imaging. The genotype of typical experimental animals has three GAL4-responsive transgenes for co-expressing proteins *A, B* and *C* labeled with eGFP, mCherry and NirFP, respectively, allowing for visualization of co-localizing POIs (FIG3d). Similarly, when a protein couple is present, the two may exhibit specific affinity with each other and can be directly demonstrated through FRET (FIG3e). Nevertheless, all FRET quantifications are to be compared to “*blank*” controls in which the fluorescently labeled POI is co-expressed with un-fused (cytoplasmic) fluorescent labels.

**Figure 3.**
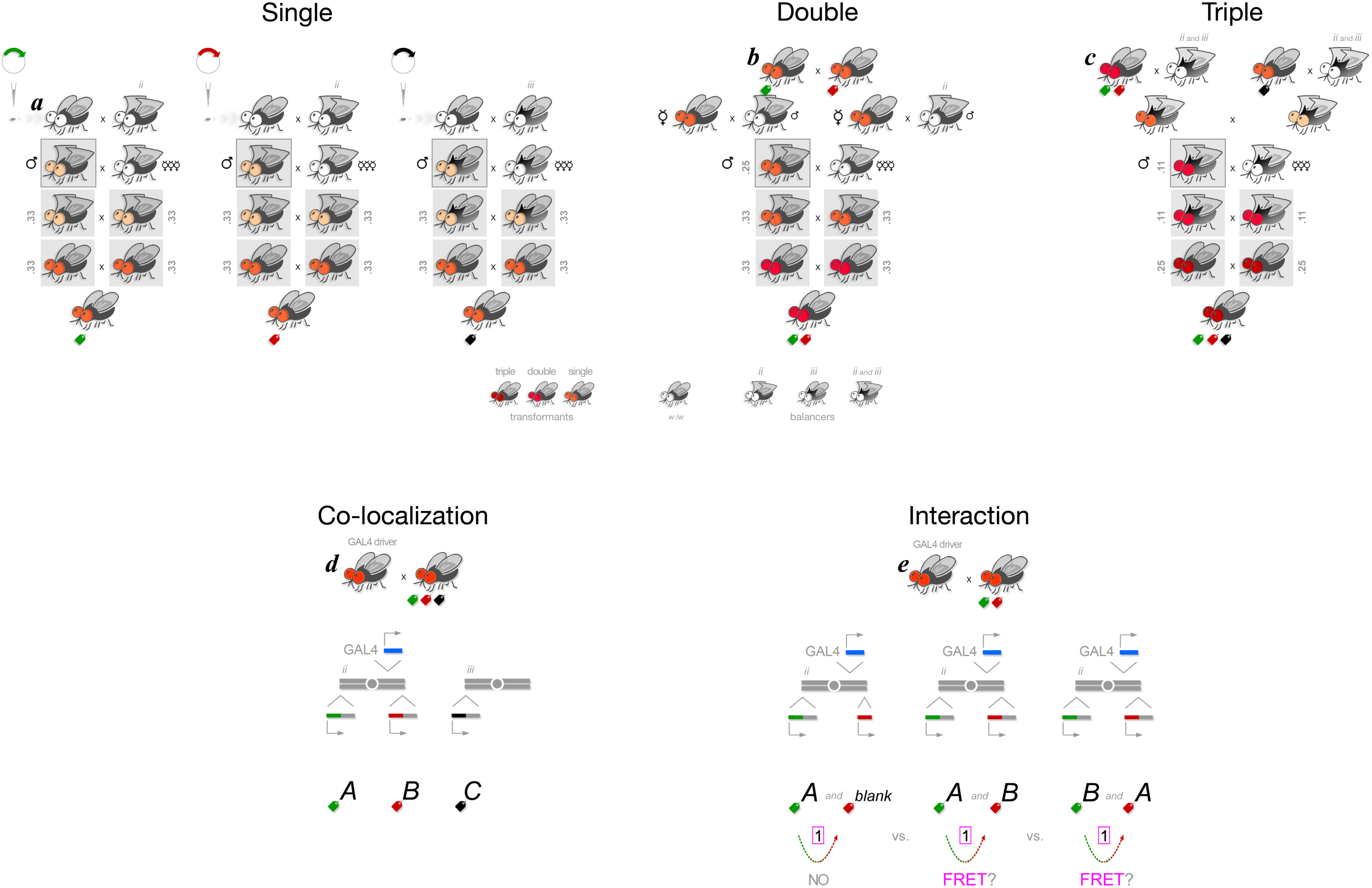
*Drosophila* genetics. (***a***) Single transgene stocks express proteins that are eGFP-labeled (*left*), mCherry-labeled (*center*) or NirFP-labeled (*right*). Five generations will: (1) integrate fluorescently labeled protein of interest (see FIG2*a*), (2) recover transformant as a single heterozygous male, (3) expand, (4) homozygose, then (5) stock. The PhiC31-mediated genomic integration into *Drosophila* genome (see FIG2*b*) occurs at a rate of approximately 2% in each sex. Generations (2) through (4) establish stock genotype with 100% confidence. When a given cross results in mixed genotypes, only those with certain genotypes are used (gray *boxes*). Probability expected using Mendelian genetics (gray *number*) and externally visible markers such as the color of eye and the shape of wing and/or hair on back cuticle are indicated. (***b***) Double transgene stocks co-express pairs of proteins that are eGFP-labeled and mCherry-labeled. Six generations will: (1) combine two transgenes on chromosome *ii* through homologous recombination, (2) cross several single females in parallel, (3) recover transformant as a single heterozygous male, (4) expand, (5) homozygose, then (6) stock. Homologous recombinations occur in the germ line of females, but balancer chromosomes suppress any such occurrence. If a virgin female mates with a male carrying the balancer for chromosome *ii*, the chance of having two transgenes on this chromosome homologously recombine with each other will be virtually 50:50. The expected offspring ratio in generation (3) will be either 50% red and 50% white in eye color with or without the balancer, or alternatively all barely red regardless of their wing being curly or not. (***c***) Triple transgene stocks co-express trios of proteins that are eGFP-labeled, mCherry-labeled and NirFP-labeled. Six generations will: (1) re-balance a double transgene stock on chromosome *ii* (*left*) and a single transgene stock on chromosome *iii* (*right*), (2) combine the three transgenes while suppressing any homologous recombination on either chromosome, (3) recover transformant as a single male, (4) expand, (5) homozygose, then (6) stock. Similar six generations of crosses starting with a single transgene stock of mCherry-labeled protein and a single transgene stock of NirFP-labeled protein establish a double transgene stock (not shown). All animals are of *w^-^/w^-^* background and, therefore, their eye colors reflect the number of transgenes they carry. Balancer for chromosome 2 (*ii*) carries *Cy^’^* mutation that causes wings to curl, and chromosome 3 (*iii*) carries *Sb^’^* mutation that causes hair on back cuticle to thicken. (***d***) Co-localization of proteins *A, B* and *C*. Three proteins labeled with eGFP, mCherry and NirFP, respectively, under the control of a GAL4 driver in embryos/larvae may be imaged by crossing fathers carrying the GAL4 driver to mothers from a triple transgene stock (see *(c)*). (***e***) Possible interaction between proteins *A* and *B*. Embryos/larvae may be imaged by crossing fathers carrying the GAL4 driver to mothers from a double transgene stock for labeling proteins with eGFP and mCherry (see *(b)*). FRET between eGFP and mCherry are considered evidence for physical interaction between the proteins *A* and *B*. Such an interaction possibility can be further confirmed through reciprocal labeling for this protein pair. Negative control (*no FRET*) to be run in parallel is eGFP-labeled protein *A* expressed along with “*blank*” cytoplasmic mCherry.

A label may be added to either the amino or carboxyl terminus of any POI, offering independent ways to visualize the same protein in the same microenvironment (FIG4a). The same protein may be labeled by any of the three colors (FIG4b). In our experience, choice in the label position or label color has not altered the intensity or distribution pattern of fluorescence, supporting their interchangeability as labels. The GAL4 driver may be ubiquitously expressed (*e.g., tub-GAL4* drives expression in all cells throughout development), specific to a whole tissue (*e.g., elav-GAL4* drives expression in all neuronal but not glia cells) or to a single cell (*e.g., eve-* GAL4 drives expression in aCC motoneurons in addition to frequent pCC interneurons and RP2 motoneurons in all 14 segments of the ventral nerve cord) (FIG4c). In neuroscience, the question of presynaptic versus postsynaptic components of synapses is important as discussed on long-term potentiation, for example (Sanes and Lichtman, 1999). Being able to limit expression to either the presynaptic neurons, postsynaptic neurons or target cells would be valuable. In a general sense, biology of multi-compartmental cells can be scrutinized in ways not easily possible with endogenous or other non-GAL4 controls. The three fluorescent colors may label three different proteins *A, B* and *C* in the same cell or cell type defined by a particular GAL4 driver (FIG5). There, proteins may be observed in patterns that are unique to each tissue and/or a single cell within a given animal. Co-expression is necessary but not predictive of their interaction, nevertheless. In fact, patterns of expression for each protein as well as those of co-expression may not correlate at all to when and where FRET occurs (FIG6). Therefore, the true protein interactomic map awaits future characterization though direct imaging of fluorescently labeled proteins within animals, *i.e.,* the natural context.

**Figure 4.**
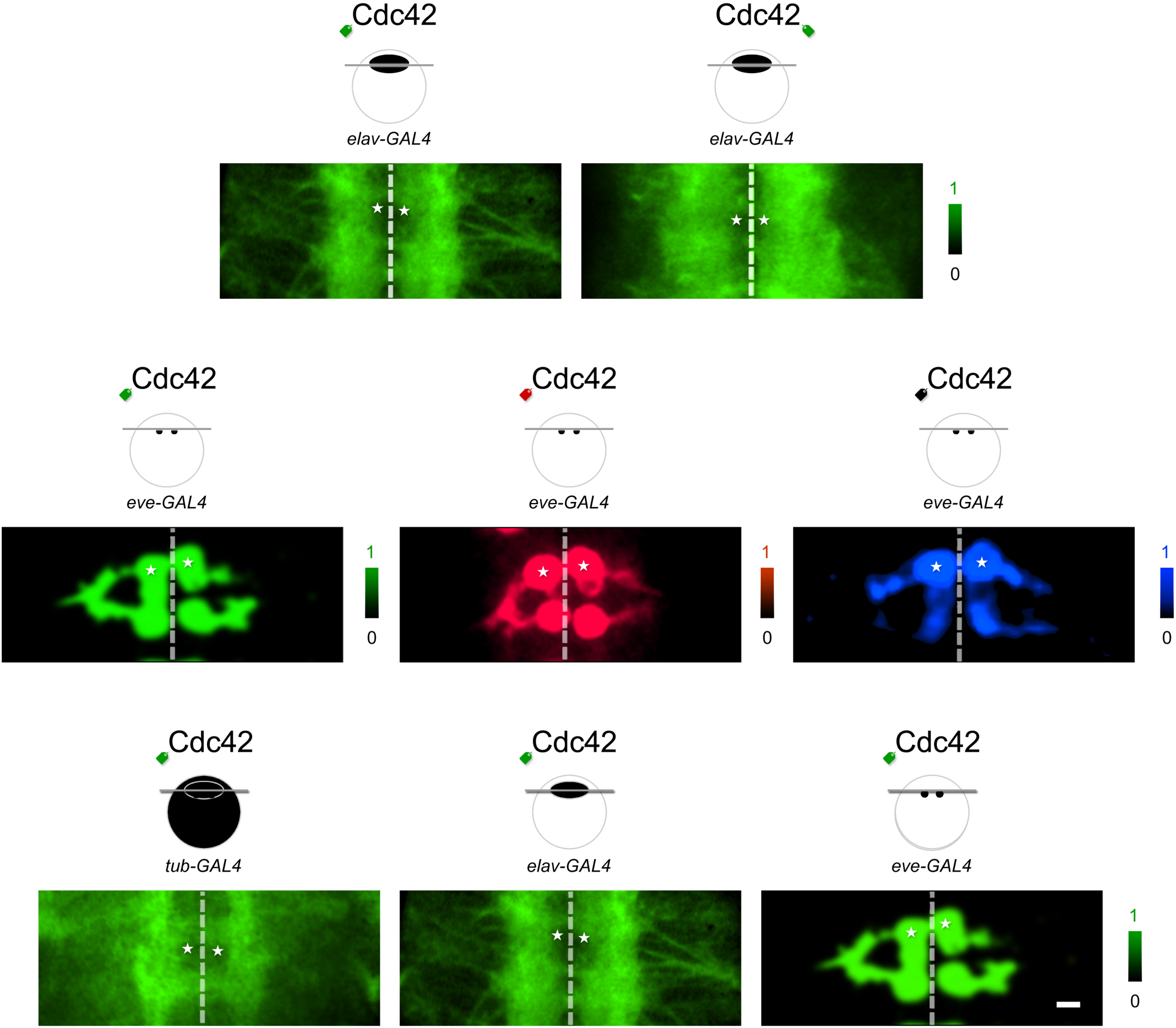
Expression control. (***a***) Label may be fused to either amino or carboxyl terminus of protein of interest. In our experience, both the amino or carboxyl termini labeling were indistinguishable. Genotypes: *UAS-eGFP::Cdc42 / elav-GAL4* (*left*), *UAS-Cdc42::eGFP / elav-GAL4* (*right*). (***b***) Label may adopt either a green color with eGFP (*left*), a red color with mCherry (*middle*) or a black (purple) with NirFP (*right*). Genotypes: *UAS-eGFP::Cdc42 / eve-GAL4* (*left*), *UAS-mCherry::Cdc42 / eve-GAL4* (*middle*), *+* / *eve-GAL4; UAS-NirFP::Cdc42 /* + (*right*). Again, colors do not seem to impact the way proteins distribute within the animals. (***c***) Labeled proteins may be expressed either ubiquitously using *tub-GAL4* (*left*), pan-neuronally using *elav-GAL4* (*middle*) or cell specifically using *eve-GAL4* (*right*). Genotypes: *UAS-eGFP::Cdc42 / tub-GAL4* (*left*), *UAS-eGFP::Cdc42 / elav-GAL4* (*middle*), *UAS-eGFP::Cdc42 / eve-GAL4* (*right*). Dashed lines indicate the midline. Asterisks point to the cell bodies of the bilateral pair of aCC motoneurons in the abdominal segment 3. Scale bar 10 nm.

**Figure 5.**
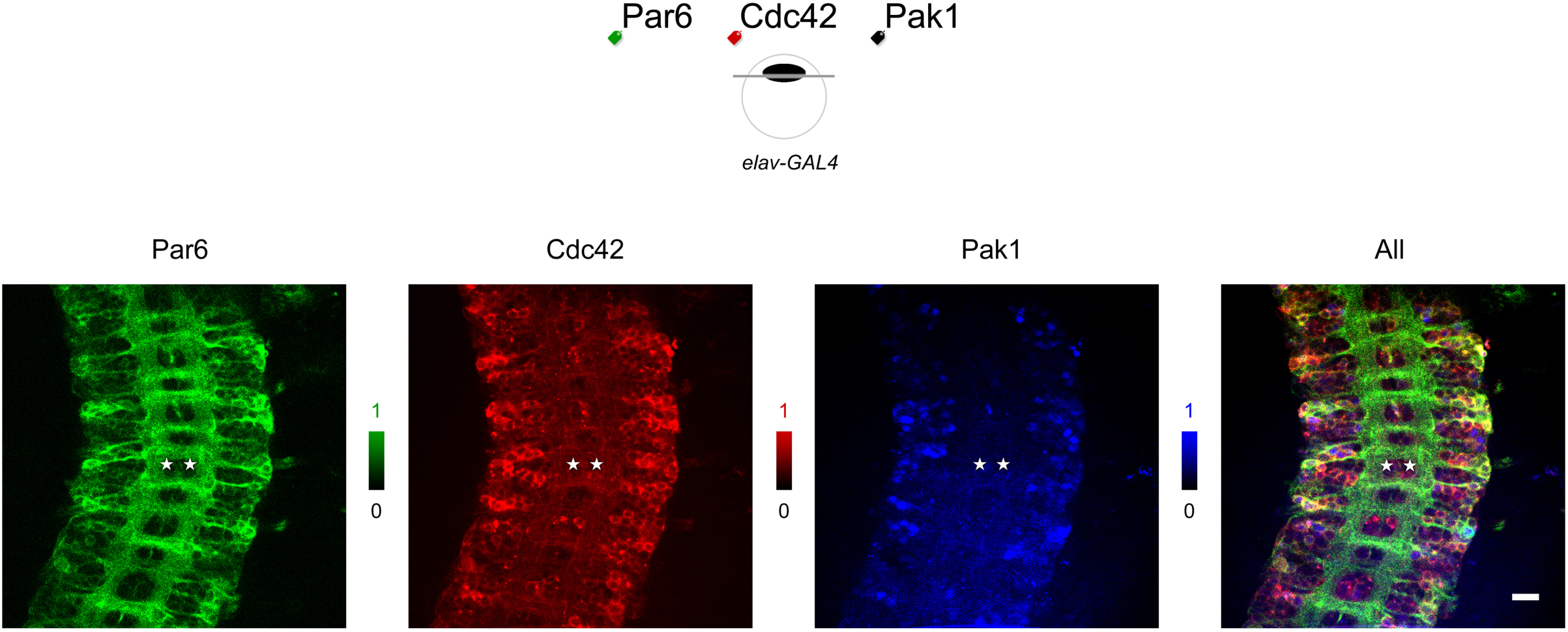
Co-localization of *A, B* and C. eGFP-labeled Par6 (green), mCherry-labeled Cdc42 (red), and NirFP-labeled Pak1 (blue *pixels*) are co-expressed in all neurons of a *Drosophila* embryo. Their expression patterns vary from pixel to pixel and from cell to cell. Anatomical features help identify a bilateral pair of aCC motoneurons (*asterisks*) in the abdominal segment A2. Genotype: *UAS-eGFP::Par6 UAS-mCherry::Cdc42 / elav-GAL4; UAS-NirFP::Pak1 / +.* Scale bar 20 nm.

**Figure 6.**
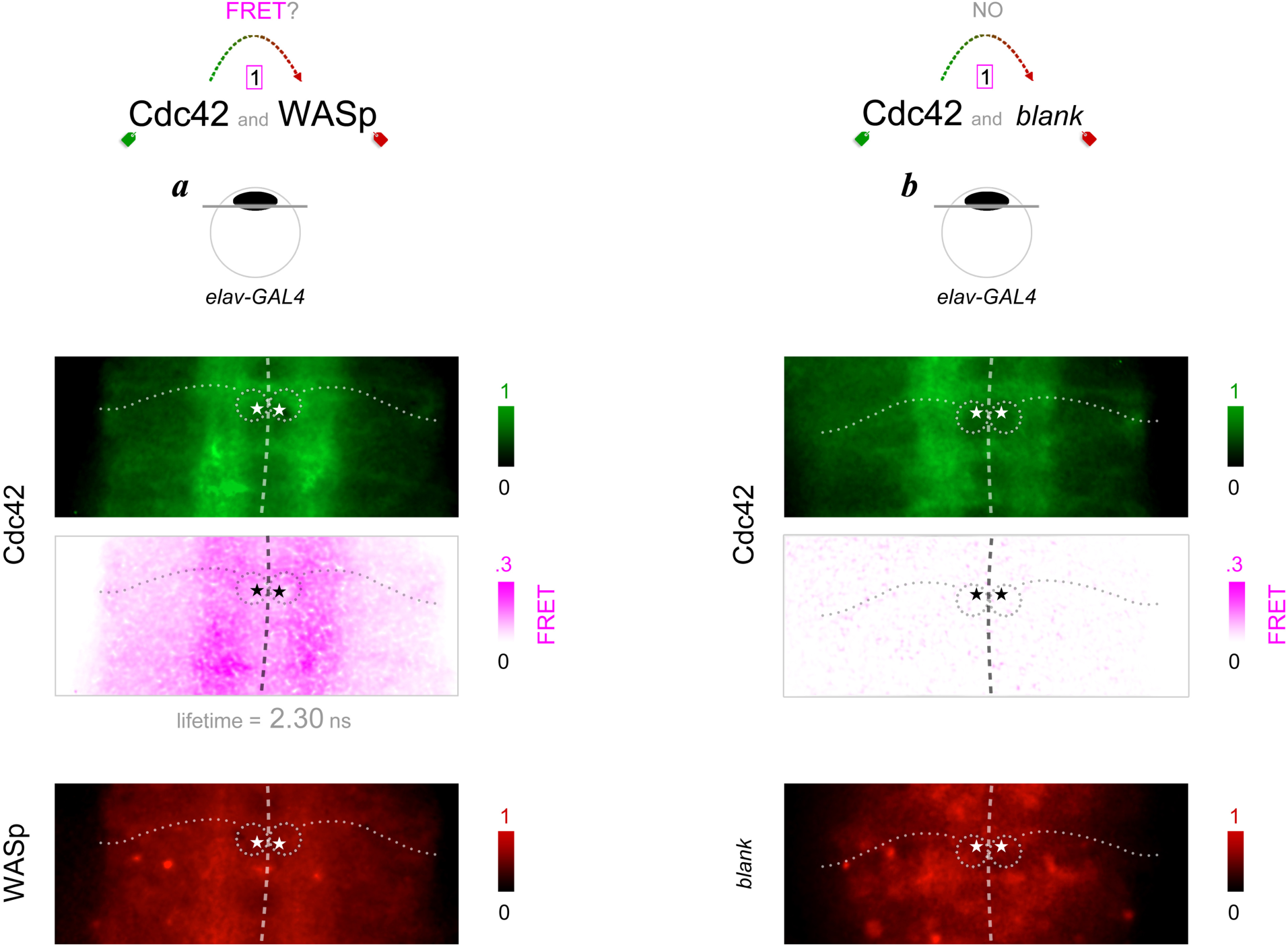
Interaction between *A* and *B.* (***a***) Interaction between monomeric G protein Cdc42 and its alleged partner WASp is revealed by FRET. Cdc42 is present at higher concentrations in neuropil region near the midline (dashed line) compared to cortical region (*top* panel). Probability of FRET between eGFP and mCherry that labels Cdc42 and WASp, respectively, is higher in the neuropil (*middle* panel). Estimated FRET probability of .24 here translates to about one in four eGFP-labeled Cdc42 proteins are within the FRET-able proximity to mCherry-labeled WASp proteins. *Asterisks* show where the bilateral pair of aCC motoneuron cell bodies are located. Their axons (dotted lines) extend laterally through first the neuropil region with higher FRET and then out into the cortical region with lower FRET. Distribution pattern of WASp, meanwhile, is distinct from Cdc42’s (*bottom* panel). However, the expression patterns of Cdc42 and WASp as well as their co-expression do not correlate to the observed FRET pattern. (***b***) Control intended for no FRET detection. The mean eGFP fluorescence lifetime at the neuropil stays at 2.56 ns, indicating no FRET. Genotype: *UAS-eGFP::Cdc42 UAS-WASp::mCherry / elav-GAL4* in (*a*) and *UAS-eGFP::Cdc42 UAS-mCherry / elav-GAL4* in (*b*).

## DISCUSSION

The labeling method described in this paper employs the exogenous GAL4 system. Potential drawbacks of this, or any other methods not using the endogenous expression control, are two-fold. First, the timing and dosage of expression can be inappropriate for assessing the function of the protein of interest. The endogenous expression patterns are often available through mRNA *in situ* hybridization and/or antibody staining of their gene products. In theory, one can focus only on sites and times that are justifiable. Even so, any difference between the native endogenous expression and controlled exogenous expression can be a cause of concerns. Second, the endogenous gene expression remains untouched. Consequently, labeled protein *A* would potentially interact with both exogenously expressed and endogenously present labeled protein *B* and un-labeled protein *B*, respectively. When looking for evidence of FRET, there is always a possibility of under-estimating the true interaction probability between proteins *A* and *B*. (*Note:* Protein trapping, that in theory circumvent issues arising from endogenous versus exogenous expression controls, has been designed for the *Drosophila* studies using green and red fluorescent proteins (Quinones-Coello et al., 2007; Venken et al., 2011); but the method described in this paper integrates notable advantages — see below.)

Our method incorporates several useful design concepts (see Table 1). In particular, the use of genetically encoded fluorescent proteins guarantees a one-to-one relationship between the label and the labeled. Also, their sturdy yet flexible linkers are thought to drastically simplify FRET computation. In addition, the GAL4 expression system, despite the discussion above, offers an unparalleled control over the time and place of expression, as well as maintaining it at a constantly low dosage. Furthermore, with position-specific transgene integration, the variability of transgene expression levels — a major source of complication in many genetic experiments — becomes minimal if not a non-issue in practice. As an added advantage, it is relatively straightforward to combine multiple labeled proteins through a series of genetic crosses. An additional detail not to be overlooked is that with an ever-expanding list of available GAL4 drivers (Jenett et al., 2012), the access to single cells becomes increasingly feasible. This would grant a significant advantage to neuroscience where axons, dendrites and synapses of the same cell can manifest distinct properties. Altogether, co-localization of three proteins *A, B* and *C,* as well as possible interaction between *A* and *B,* independent of the interaction between *B* and *C,* may all be documented. Finally, principles from most of these genetic engineering tools may be applicable to other metazoan model organisms such as nematodes, zebra fish and small mammals. Whether or not the concepts used in design of labeling or expression control may vary in detail, proteins that are labeled fluorescently in diverse animal systems will open a way to map freely interacting proteins within their natural context.

## METHODS

### Acceptor Vector construction

To generate the C-terminus tagged vectors, the pUAST expression vector (Brand and Perrimon, 1993) was cut with NotI and XbaI. The pUAST-C-TAP-BD expression vector (Stapleton) was cut with NotI and XbaI to release a 256 bp fragment containing loxP, chloramphenicol promoter and splice acceptor sequences. This 256 bp C-terminus BD cassette was inserted into the linearized pUAST vector by overnight ligation. The resulting C-terminus BD-adapted pUAST vector was linearized again by cutting it with XbaI. To generate the N-terminus tagged vectors, the pUAST expression vector was cut with EcoRI and XhoI. The pUAST-N-TAP-BD expression vector was cut with EcoRI and XhoI to release a 167 bp fragment containing loxP and chloramphenicol promoter sequences. This 167 bp C-terminus BD cassette was inserted into the linearized pUAST vector by overnight ligation. The resulting N-terminus BD-adapted pUAST vector was linearized again by cutting it with EcoRI. The CDS of the EGFP and mCherry fluorescent tags were obtained by PCR from the pEGFP-C1 (Clontech) and pRSET-B-mCherry (Invitrogen) vectors, respectively. C-terminus fluorescent tags were amplified using XbaI-tailed PCR primers, while N-terminus tags were amplified using EcoRI-tailed PCR primers (see below). The resulting PCR products were agrose gel-purified and cut with either XbaI (C-terminus tag) or EcoRI (N-terminus tag). Following ethanol precipitation, the XbaI digested C-terminus fluorescent tags were inserted into the XbaI linearized BD-adapted pUAST vector by overnight ligation. The EcoRI digested N-terminus tags were inserted into the EcoRI linearized BD-adapted pUAST vector by overnight ligation. The resulting C- or N-terminus fluorescent-tagged and BD-adapted pUAST vectors are cut with BamHI to release the portion of the vector containing the 5x UAS promoter, hsp70, BD cassette, fluorescent tag and SV40 terminator sequences. Agarose gel-purify the BamHI digested fragments. We linearized the pBDP vector by cutting it with BamHI and ethanol precipitated the digested product. We inserted the purified C- or N-terminus fluorescent-tagged and BD-adapted BamHI fragments from the pUAST vectors into the linearized pBDP vector by overnight ligation. The resulting vectors are pUAS-C-EGFP-BD-attB, pUAS-C-mCherry-BD-attB, pUAS-N-EGFP-BD-attB and pUAS-N-mCherry-BD-attB. (*Note:* Expression vectors were designed similarly for labeling POI with NirFP but are yet unavailable at the DGRC.)

Sequences: 256bp C-terminus BD cassette: 5’-GGCCGCATAACTTCGTATAGCATACATTATACGAAGTTATAGATCCAATATTATTGAAGCAT TTATCAGGGTTATTGTCTCATGAGCGGATACATATTTGAATGTATTTAGAAAAATAAACAAAT AGGGGTTCCGCGCACATTTCCCCGAAAAGTGCCACCTGACGTGGATCTCGAGCTCAAGCT TCGAATTCAGGGTTTCCTTGACAATATCATACTTATCCTGTCCCTTTTTTTTCCACAGCTACC GGTCGCGT-3’ 167bp N-terminus BD cassette: 5’-AATTCATAACTTCGTATAGCATACATTATACGAAGTTATAGATCCAATATTATTGAAGCATTT ATCAGGGTTATTGTCTCATGAGCGGATACATATTTGAATGTATTTAGAAAAATAAACAAATA GGGGTTCCGCGCACATTTCCCCGAAAAGTGCCACCTGACGTC-3’ XbaI-tailed PCR primers: C-mCherry-FWD 5’-cccctctagaGTGAGCAAGGGCGAGGAGGATAACATG-3’ C-mCherry-REV 5’-ggggtctagaTTACTTGTACAGCTCGTCCATGCCGC-3’ C-EGFP-FWD 5’-cccctctagaGTGAGCAAGGGCGAGGAGCTGTTC-3’ C-EGFP-REV 5’-ggggtctagaTTACTTGTACAGCTCGTCCATGCCGAG-3’ EcoRI-tailed PCR primers: mCherry-N-FWD 5’-ccccgaattcacaccATGGTGAGCAAGGGCGAGGAGGAT-3’ mCherry-N-REV 5’-ggggaattcccCTTGTACAGCTCGTCCATGCCGCC-3’ EGFP-N-FWD 5’-ccccgaattcacaccATGGTGAGCAAGGGCGAGGAGCTG-3’ EGFP-N-REV 5’-ggggaattcccCTTGTACAGCTCGTCCATGCCGAG-3’

### Expression Clone sets

We transferred ORFs from the BDGP *Drosophila melanogaster* XO and XS expression-ready clone sets to the pUAS-C-EGFP-BD-attB, pUAS-C-mCherry-BD-attB, pUAS-N-EGFP-BD-attB and pUAS-N-mCherry-BD-attB acceptor vectors (Yu et al., 2011). For recombination reactions, 200 ng of expression-ready Donor clone and acceptor Vector were recombined in a final volume of 10 µl for 15 minutes at 25 °C in a thermal cycler in the presence of Cre recombinase (0.2 units) and recombinase buffer supplemented with BSA (0.1 mg/ml) (Clontech #631614) according to Clontech manual PT3460-1 (now part of Takara Bio). Cre recombinase was inactivated by incubating the reaction at 70 °C for 10 minutes. From this reaction, 5 µl was transformed into chemically competent TAM-1 cells in a 96-well plate format (Active Motif #11096) and selected for Chloramphenicol resistance. Each clone was sequence verified to check for target mismatches using BigDye Terminator v3.1 ready reaction mix (Applied Biosystems #4337457) and the sequencing primer: 5’-GCCAATGTGCATCAGTTGTGGTC-3’. Sequencing samples were analyzed on a conventional capillary electrophoresis instrument (e.g., ABI 3730/3730xl DNA Analyzer). Glycerol stocks were generated and stored for each isolate. Clones can be obtained from the *Drosophila* Genome Resource Center (DGRC) in Bloomington, IN and sequence from GenBank using the accession numbers provided in Supplemental Table.

Drosophila *stocks. tub-*GAL4 - source: Bloomington Stock #60298 was used for ubiquitous expression. *elav-*GAL4 - source: Bloomington Stock # 8760 was used for pan-neuronal expression. *eve-*GAL4[*^RN2-^*^GAL4*-E*^] - source: M. Fuijoka was used for expression in aCC/RP2/pCC neurons. Transgenic flies carrying *UAS-eGFP::POI* (*attp40* site at 25C7 - source: www.geneticservices.com), *UAS-mCherry:POI* (*vk01* site at 59D3 - source: Bloomington Stock # 9722) on the second chromosome and *UAS-NirFP::POI* (*attp2* site at 68A4 - source: Bloomington Stock # 8622) on the third chromosome were constructed as described (Sharifai et al., 2014). All *attp* landing sites were selected because of their integration rates varying as high as 25-35%, *i.e.,* more efficiently than *P* element transformation of about 1%. *P Bac(y+.attp-3b) attp2* and *P Bac(y+.attp-3b) att40* are from the collections described in (Groth et al., 2004) and (Szabad et al., 2012). They are located at map positions 25C6 (*attp40*) on the second chromosome and 68A4 (*attp2*) on the third chromosome. *PBac(y+.attp-3b) vk01* is from the Venken Bac collection (Venken et al., 2006b) and is located at 59D3 of the second chromosome. Besides their high integration rates, these integration sites were used also for high expression rates and low background effects (Markstein et al., 2008; Pfeiffer et al., 2010) (Zusman et al. unpublished observations). For all integration sites the integrase-producing transgene *P(nos-phiC31\int.NLS) X* (Bischof et al., 2007) was used.

### Imaging

UAS-transgene carrying males were crossed with GAL4 driver carrying virgin females in mating cages using grape juice agar plates coated with yeast to collect embryos. Mating cages were placed at 25 °C and collection plates were swapped after one hour to gather embryos for a given developmental stage. Embryos were manually dechorionated and developmental staging was confirmed by gut morphology. Embryos were placed on double-sided tape attached to a glass slide with a silicone well and then immersed in HL3.1 hemolymph buffer for imaging. We used an upright microscope with a x40 or x63 water-immersion objective lens to image the sample. However, when an inverted microscope was used, a coverslip was placed over the buffer-immersed embryos using elastic silicone. Subsequently, the slide was flipped upside-down and mounted on a x40 or x63 oil-immersion objective lens. In some cases, devitillenization, filet dissection, or both, were performed to increase signal intensities of eGFP, mCherry and NirFP. A ratio of mCherry over eGFP being between 0.1 and 10 was found essential for obtaining reliable FRET measurements, consistent with previous findings (Berney and Danuser, 2003). No samples created through our transgenic method produced a sign of “false positive” FRET, *i.e.,* a false decrease in donor fluorescence lifetime. In contrast, experiments using standard retrovirus transfection methods in human and insect cell lines had an expression level of mCherry, the FRET acceptor, beyond the ratio of 100 and produced such a “false positive” FRET (not shown).

### FRET probability

The probability of FRET is defined as a product of the concentration and lifetime reduction of a FRET donor, *e.g.,* eGFP or mCherry. Whereas the concentration of a given protein varies from pixel to pixel, the fluorescence lifetime *τ* is calculated independently of its concentration. Consequently, a polar histogram allows for normalization of the FRET probability irrespective of the size of proteins interacting with each other. This size-free unit of protein interactome is termed “interaction” (Sharifai et al., 2014).

## Acknowledgements

We thank TK Harris and Rajeev Probhakar for structural biology of GFP, Konstantin Lukyanov for chemistry of NirFP, Edward Giniger, Francisco Raymo, and Peter Larsson for discussions on FRET-based protein imaging, Grace Zhai for discussion on Drosophila, Laura Bianchhi and Kevin Collins for discussion on C. elegans, Julia Dallman and Sandra Rieger for discussion on zebra fish, and Vance Lemmon, Patelis Tsoulfas and Tom Lisse for discussion on mice.

Supported in part by research awards RC2-NS069488 from American Recovery and Reinvestment Act and National Institutes of Health and MH079432 from National Institutes of Health to AC, and research award P41HG3487 from National Human Genome Research Institute to SC, and performed under U.S. Department of Energy Contracts DE-AC0376SF00098 and DE-AC02-05CH11231.

---

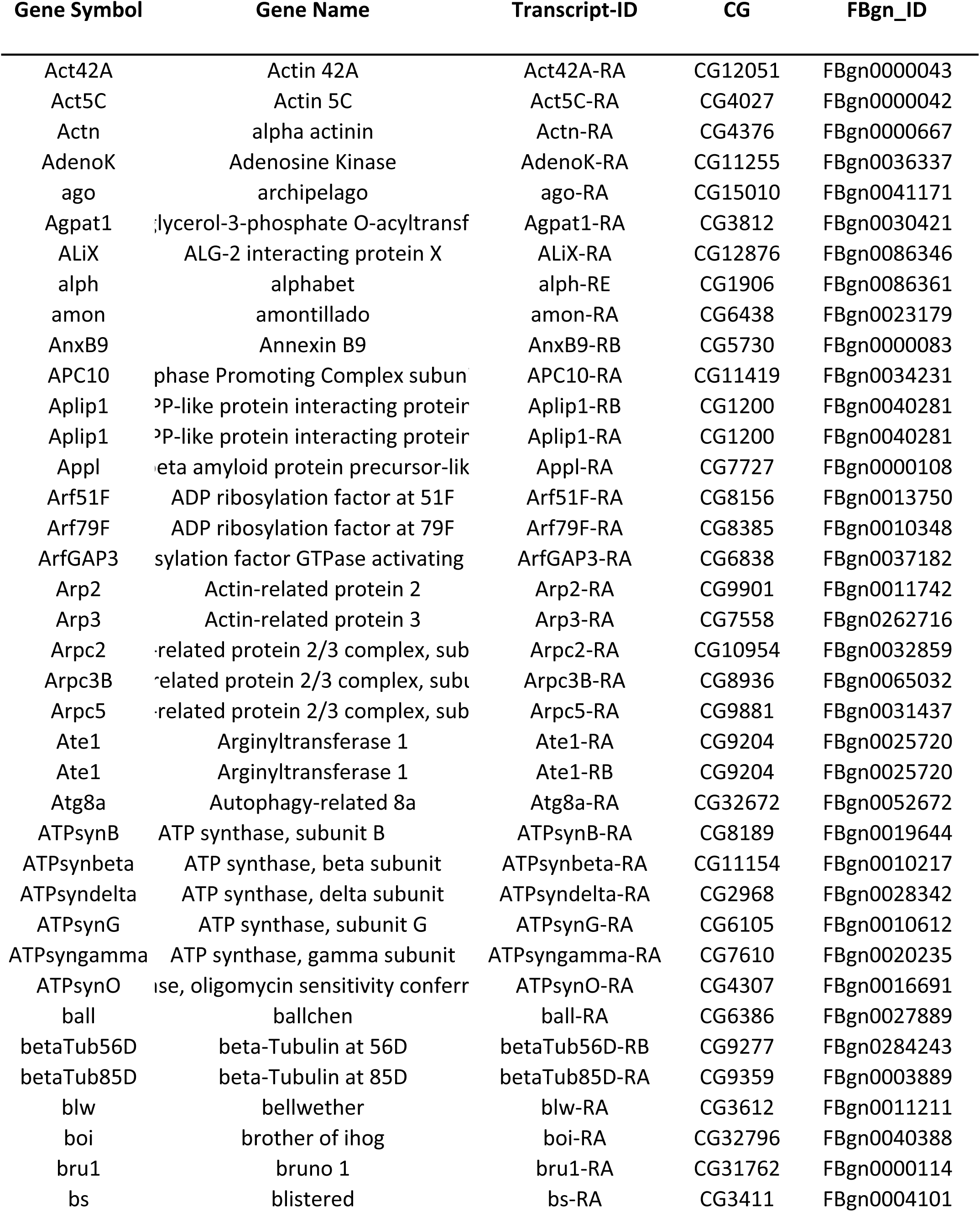
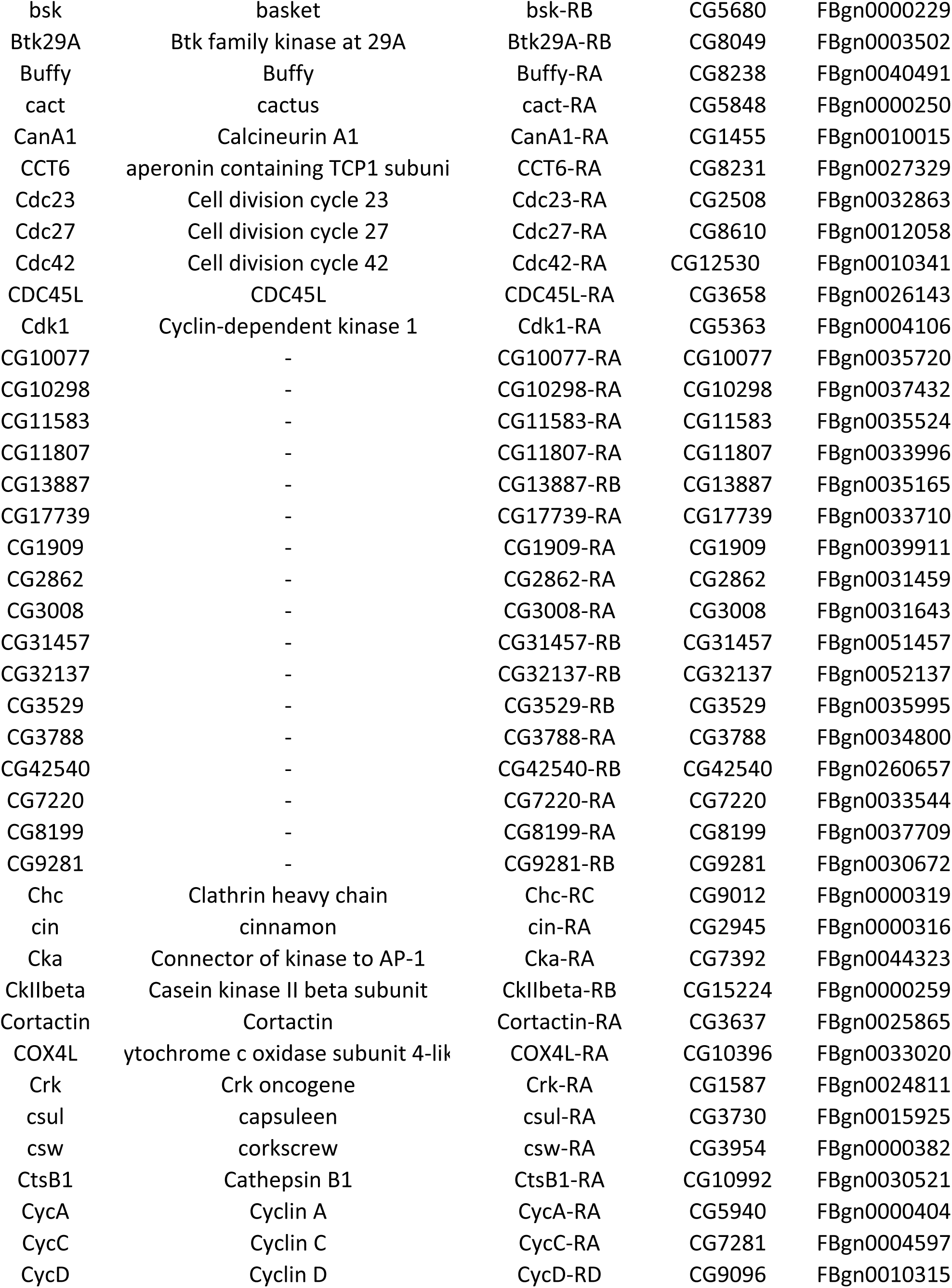
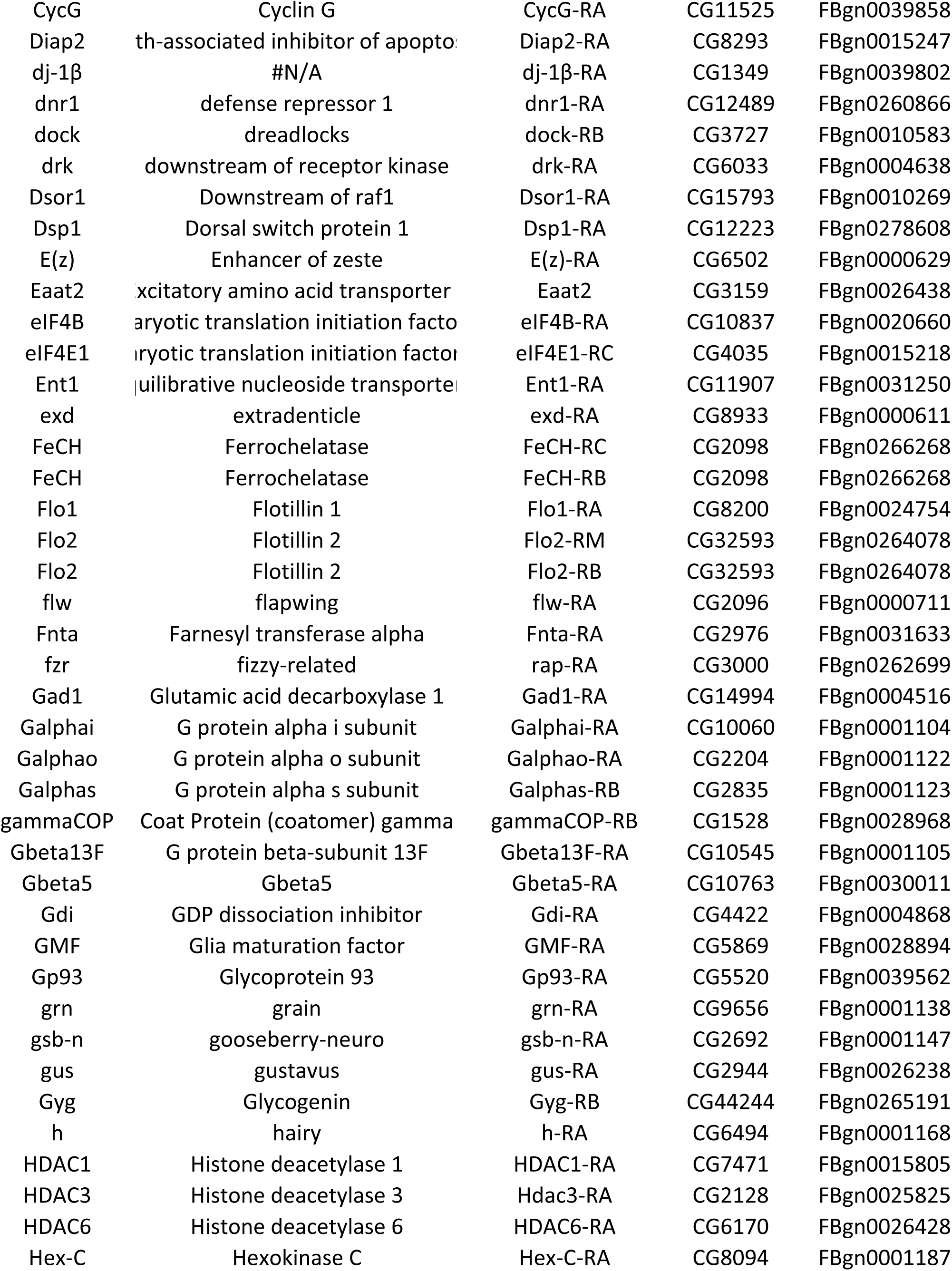
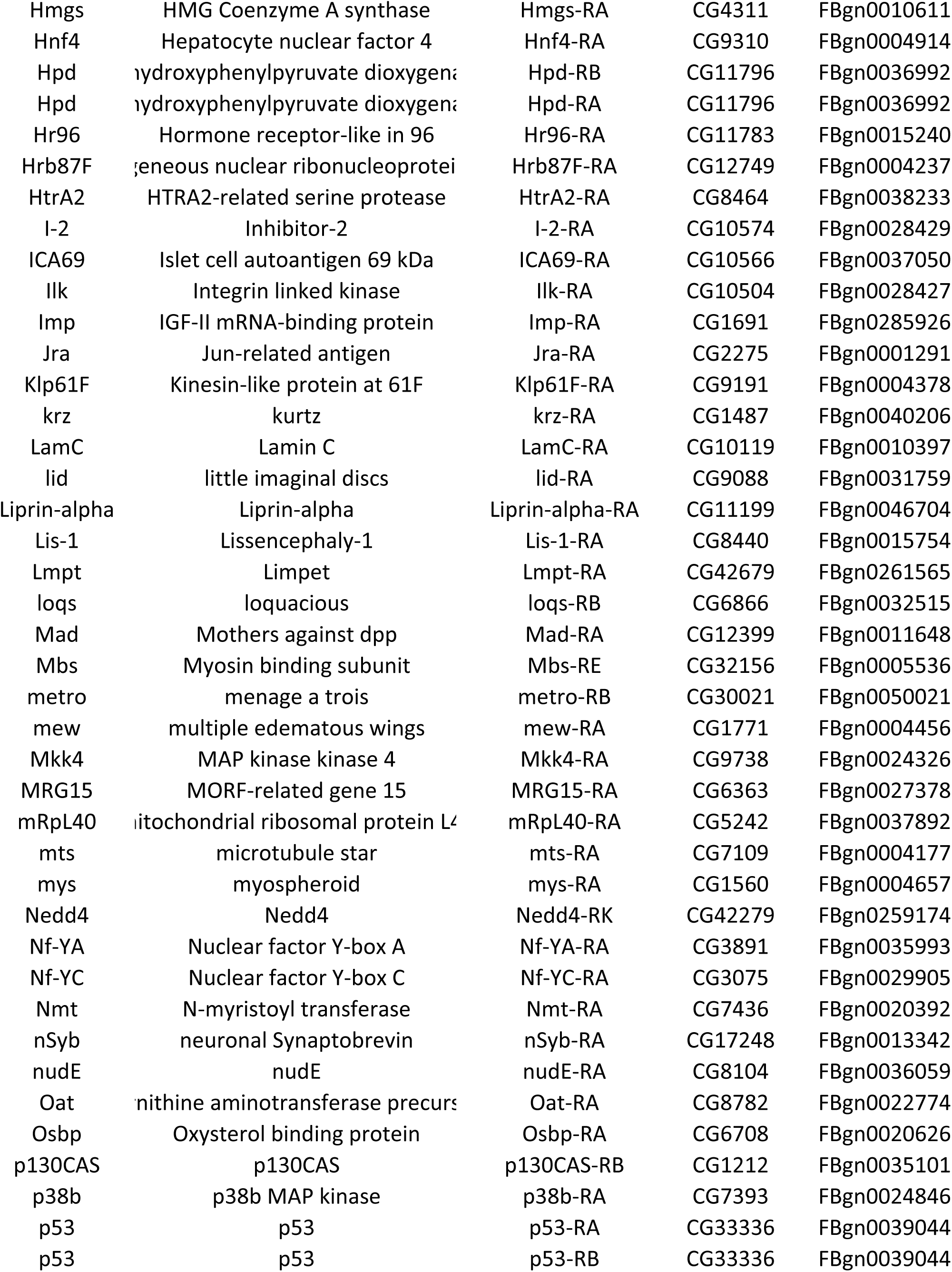
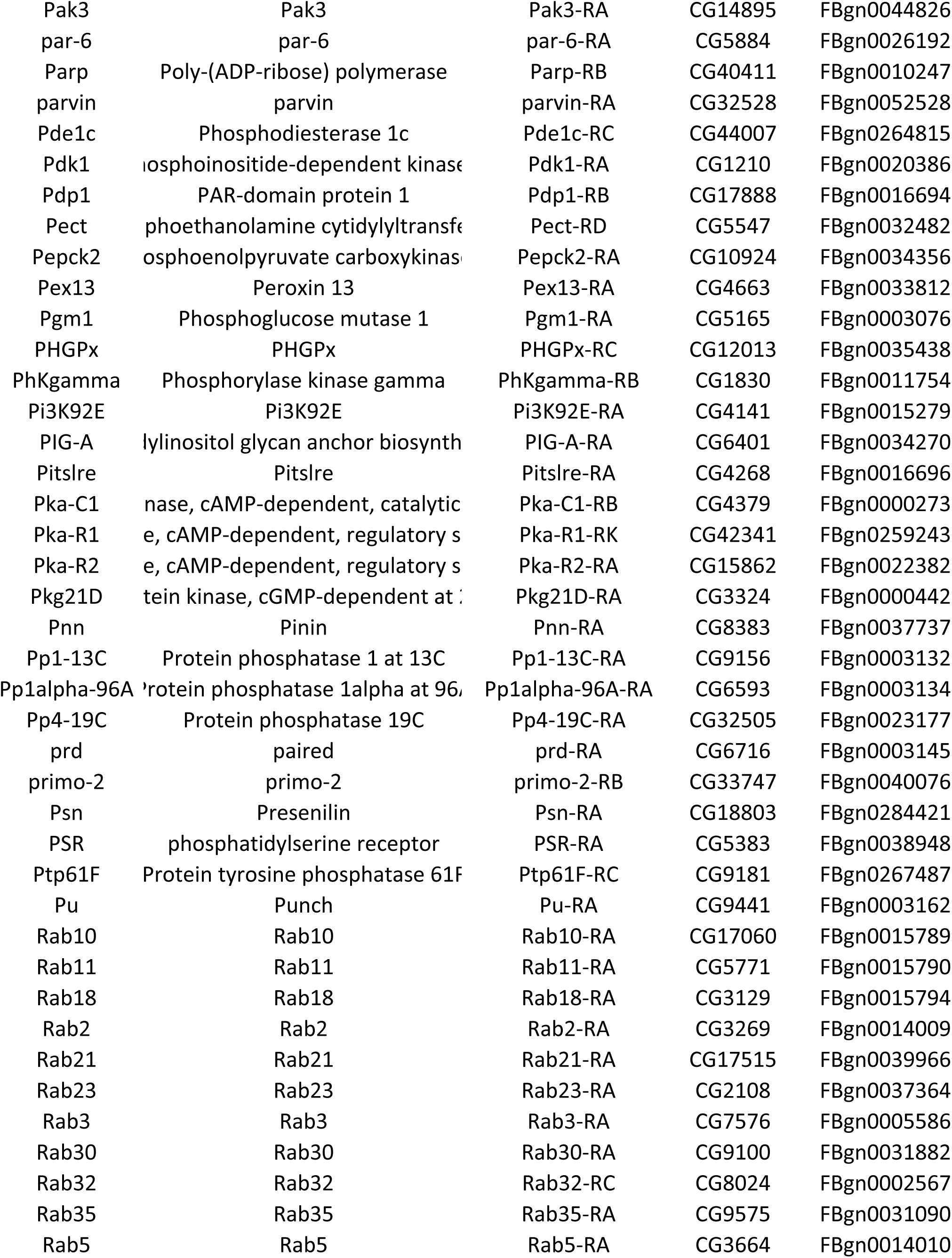
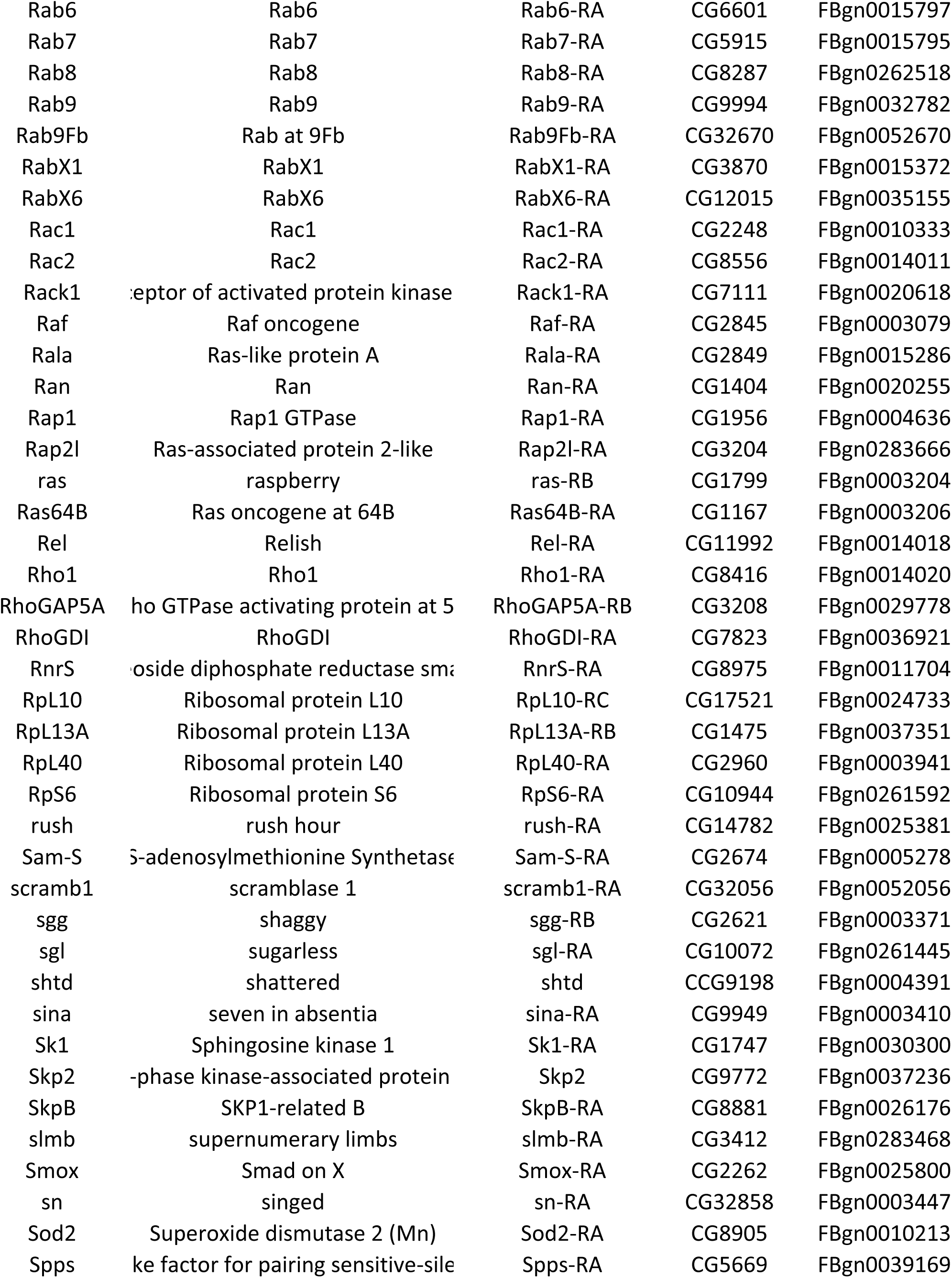
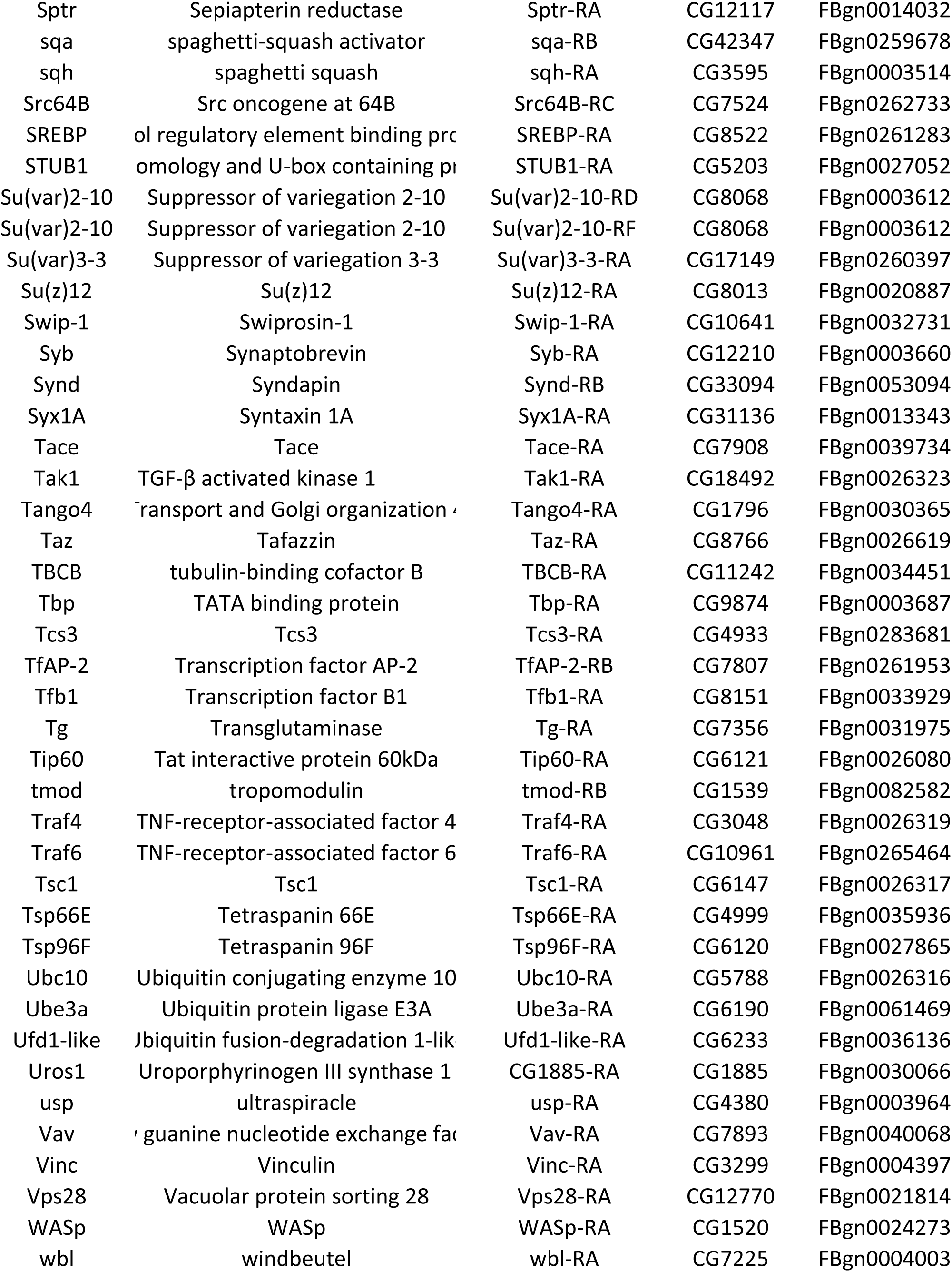
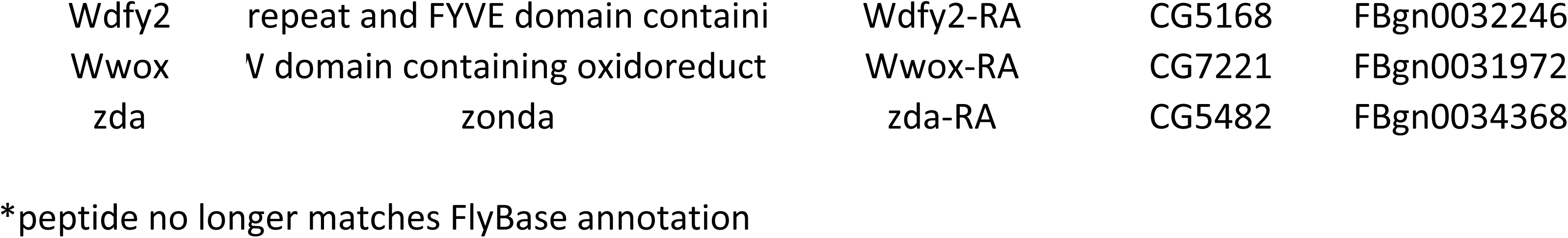

| CDS size | Gold Source Clone | Accession number | Donor clone stop (DS) | DS Accession number | GFP::ORF | RFP::ORF |
| --- | --- | --- | --- | --- | --- | --- |
| 1128 | LD18090 | AY118907 | BS15418 | KX801817 | UGS01407 | URS01407 |
| 1128 | RE02927 | AY089562 | BS11334 | FJ636827 | UGS01271 | URS01271 |
| 2685 | LD37956 | AY069615 | ND | ~ |  |  |
| 1035 | GH14845 | AY060709 | BS09556 | FJ638826 | UGS01220 | URS01220 |
| 3978 | LD21322 | AY061300 | BS11048 | KX802002 | UGS01492 | URS01492 |
| 1029 | LD44987 | AY118588 | BS08872 | MN238604 | UGS01215 |  |
| 2508 | LD25543 | AY069534 | ND | ~ |  |  |
| 1122 | LD23542 | AY051685 | BS06802 | FJ635621 | UGS01372 | URS01372 |
| 1962 | GH12584 | AY119509 | ND | ~ |  |  |
| 972 | LD09947 | AY118504 | BS04048 | FJ634842 | UGS01331 | URS01331 |
| 585 | RE25242 | AY071207 | BS11859 | FJ636999 | UGS01487 | URS01487 |
| 1470 | GH14842 | AY051495 | BS07363 | FJ635831 | UGS01385 | URS01385 |
| 1449 | RH26053 | BT001795 | BS15971 | FJ638001 | UGS01415 |  |
| 2661 | GH04413 | AF181628 | ND | ~ |  |  |
| 525 | RE16882 | AY071116 | ND | ~ |  |  |
| 546 | LD24904 | AY060375 | BS05649 | FJ635191 | UGS01351 | URS01351 |
| 1656 | LD46935 | AY069700 | BS10163 | KX804690 | UGS01257 | URS01257 |
| 1182 | LD18955 | AY061256 | BS04230 | FJ634886 | UGS01357 | URS01357 |
| 1254 | LD35711 | AY051859 | BS08482 | FJ638812 | UGS01204 | URS01204 |
| 903 | LD29815 | AY060391 | BS08078 | FJ636093 | UGS01202 | URS01202 |
| 522 | SD24339 | AY119268 | BS15269 | FJ637907 | UGS01472 | URS01472 |
| 453 | RH48239 | AY089678 | BS13858 | FJ637540 | UGS01256 | URS01256 |
| 1431 | LD23642 | AY051688 | BS17605 | KX800222 | UGS01423 | URS01423 |
| 1452 | LD33682 | BT001498 | BS17738 | KX800750 | UGS01435 | URS01435 |
| 363 | LD05816 | AY122152 | BS03649 | FJ638771 | UGS01328 | URS01328 |
| 729 | RE36486 | BT011453 | BS16081 | FJ638055 | UGS01478 | URS01478 |
| 1515 | RE10864 | AY118367 | BS11538 | FJ636899 | UGS01430 | URS01430 |
| 471 | RE24457 | AY071203 | BS11948 | FJ637029 | UGS01463 | URS01463 |
| 297 | RE20862 | AY071165 | BS11796 | FJ636975 | UGS01279 | URS01279 |
| 891 | RE35715 | AY113454 | BS12071 | FJ637060 | UGS01282 | URS01282 |
| 627 | GH01760 | AY058261 | BS06972 | FJ635679 | UGS01378 | URS01378 |
| 1797 | LD09009 | AF160917 | BS03617 | FJ634726 | UGS01237 | URS01237 |
| 1341 | LD43681 | BT003242 | BS15123 | KX806269 | UGS01406 | URS01406 |
| 1338 | GH02051 | AY118725 | BS14906 | FJ637829 | UGS01404 |  |
| 1656 | HL08087 | BT003592 | BS10401 | FJ636639 | UGS01228 | URS01228 |
| 1641 | RE01910 | AY094843 | BS11320 | FJ636821 | UGS01269 | URS01269 |
| 1812 | LD29068 | AY051758 | BS08061 | FJ636084 | UGS01247 | URS01247 |
| 1347 | RE17834 | AY095071 | BS11757 | FJ636960 | UGS01315 | URS01315 |
| 1116 | HL02677 | AY070865 | BS04602 | FJ634972 | UGS01339 | URS01339 |
| 2358 | LD16208 | AY128441 | BS05076 | KX795404 |  | URS01443 |
| 897 | AT16536 | AY075219 | BS14274 | FJ637627 | UGS01295 | URS01295 |
| 1500 | LD10168 | AY069399 | BS01549 | FJ634644 | UGS01317 | URS01317 |
| 1866 | RE54552 | AY128480 | BS12368 | KX805643 | UGS01468 | URS01468 |
| 1599 | GH13725 | AY051470 | BS07223 | FJ635773 | UGS01242 | URS01242 |
| 2034 | LD09850 | BT010078 | BS05017 | KX794414 |  | URS01442 |
| 2700 | LD12661 | BT021947 | BS15481 | KX801947 | UGS01473 | URS01473 |
| 573 | HL08128 | AY119570 | BS03306 | FJ634713 |  |  |
| 1725 | LD35753 | AY051861 | BS08484 | FJ636179 | UGS01251 | URS01251 |
| 891 | LD38718 | AY061450 | BS06587 | FJ635552 | UGS01365 | URS01365 |
| 2454 | GH10652 | AY102660 | ND | ~ |  |  |
| 561 | GH28351 | AY113352 | BS14872 | FJ637815 | UGS01471 | URS01471 |
| 1068 | LD22010 | AY118916 | BS05610 | FJ635161 | UGS01350 | URS01350 |
| 1473 | SD03973 | AY052088 | BS09340 | FJ636400 | UGS01219 | URS01219 |
| 684 | LP10861 | AY069750 | BS06288 | FJ635432 | UGS01360 | URS01360 |
| 2619 | GH02025 | AY058267 | BS07277 | FJ635800 | UGS01243 | URS01243 |
| 1818 | GH22690 | AY051576 | BS07690 | FJ635951 | UGS01246 | URS01246 |
| 450 | RH49748 | AY071745 | BS13964 | FJ637568 | UGS01292 | URS01292 |
| 1809 | RE33807 | AY071312 | BS12053 | KX804479 | UGS01464 |  |
| 1011 | LD10183 | AY069400 | BS04050 | FJ634844 | UGS01332 | URS01332 |
| 1860 | RE18568 | AY095527 | BS11775 | KX805359 | UGS01462 | URS01462 |
| 1629 | GH05942 | AY058297 | BS07113 | FJ635733 | UGS01382 | URS01382 |
| 1134 | AT31332 | AY075301 | BS12731 | FJ637281 | UGS01401 | URS01401 |
| 1194 | RE02540 | BT021331 | BS16738 | FJ638186 | UGS01434 | URS01434 |
| 462 | AT19555 | AY070806 | BS12785 | FJ637292 | UGS01403 | URS01403 |
| 1317 | GH19141 | AY051542 | BS07744 | MN238600 | UGS01388 | URS01388 |
| 1833 | LD02975 | AY061059 | BS03015 | FJ634690 | UGS01239 | URS01239 |
| 5034 | LD43101 | AY119615 | BS11028 | KX802159 | UGS01486 | URS01486 |
| 1803 | GH09380 | AY069078.1 | ND | ~ |  |  |
| 2190 | LD41760 | AY061460 | ND | ~ |  |  |
| 645 | RE31047 | AY113447 | BS12026 | FJ637039 | UGS01281 | URS01281 |
| 1677 | LD29964 | AY051774 | BS08083 | MN238597 | UGS01248 | URS01248 |
| 528 | AT07685 | AY084105 | BS13760 | FJ637521 | UGS01293 | URS01293 |
| 813 | RE60886 | BT012456 | BS16314 | FJ638116 |  | URS01418 |
| 1830 | LD07634 | AY122153 | BS15459 | KX803675 | UGS01408 | URS01408 |
| 2523 | LD22829 | BT001484 | BS03210 | FJ634694 | UGS01235 | URS01235 |
| 1020 | GH06546 | AY060640 | BS07022 | FJ635692 | UGS01383 | URS01383 |
| 1473 | LD44443 | AY058712 | BS08955 | FJ636323 | UGS01212 | URS01212 |
| 801 | LD35705 | AY061425 | BS06196 | FJ635392 | UGS01358 | URS01358 |
| 1431 | LD22957 | AY069509 | BS06692 | FJ635590 |  | URS01371 |
| 1698 | LD46284 | AY058720 | BS10622 | FJ636696 |  | URS01429 |
| 1494 | LD34777 | AY051844 | BS08565 | FJ636199 |  | URS01203 |
| 564 | GH09983 | AY060670 | BS06523 | FJ635531 | UGS01362 |  |
| 2028 | LD18186 | AY095032 | BS04649 | KX794325 |  | URS01441 |
| 1644 | LD42588 | AY119614 | BS08921 | FJ636318 | UGS01209 | URS01209 |
| 633 | LD12029 | AY061142 | BS02810 | FJ634676 | UGS01238 | URS01238 |
| 1188 | LD41207 | AY058692 | BS08685 | FJ636235 | UGS01252 |  |
| 1155 | GM02110 | AY060841 | BS04728 | FJ635017 | UGS01341 | URS01341 |
| 2280 | LD30505 | AY051785 | BS01296 | FJ634590 | UGS01322 | URS01322 |
| 1116 | GH09856* | AY069083 | BS16823 | KX805932 | UGS01421 | URS01421 |
| 1170 | RE54950 | AY071474 | BS12277 | FJ637130 | UGS01326 | URS01326 |
| 744 | RE36735 | BT012467 | BS16311 | FJ638114 | UGS01417 | URS01417 |
| 1428 | LD19162 | AY061261 | BS04234 | FJ634887 | UGS01333 | URS01333 |
| 1128 | LD03509 | AY061065 | BS03889 | FJ634814 | UGS01305 | URS01305 |
| 843 | GH01314 | AY058251 | BS06964 | FJ635675 | UGS01377 | URS01377 |
| 678 | SD11336 | BT001878 | BS17974 | KX803751 | UGS01426 | URS01426 |
| 1278 | SD10657 | AY058794 | BS09502 | FJ636432 |  | URS01447 |
| 1068 | LP11503 | AY069755 | BS09979 | FJ636515 | UGS01223 | URS01223 |
| 1275 | RE74011 | BT009962 | BS16376 | MN238598 |  | URS01431 |
| 1383 | GH05039 | AY069047 | BS07406 | FJ635845 |  | URS01494 |
| 993 | LD26412 | AY118534 | BS06620 | MN238602 | UGS01366 | URS01366 |
| 1434 | GH07620 | BT001403 | BS15605 | KX802445 | UGS01411 | URS01411 |
| 1530 | HL02049 | AY089526 | BS05072 | FJ635055 | UGS01344 | URS01344 |
| 1065 | LD22201 | AY051670 | BS06675 | FJ635585 | UGS01370 | URS01370 |
| 1062 | GM01620 | AY121631 | BS04626 | MN238599 |  | URS01340 |
| 1155 | LD26182 | AY058572 | BS10509 | FJ636687 |  | URS01229 |
| 2649 | RE37840 | BT001628 | BS14096 | KX800981 | UGS01233 | URS01233 |
| 1020 | LD25526 | AY058566 | BS08015 | FJ636059 | UGS01394 |  |
| 1074 | RH17413 | BT004841 | BS13476 | FJ638854 | UGS01289 | URS01289 |
| 1329 | LP03430 | BT003563 | ND | ~ |  |  |
| 414 | RE40543 | BT014645 | BS16196 | KX803490 | UGS01477 | URS01477 |
| 2361 | LD23641 | AY075384 | BS10205 | FJ636594 | UGS01485 | URS01485 |
| 1458 | RE40104 | AY071369 | BS01543 | FJ634641 | UGS01309 | URS01309 |
| 1347 | RE64348 | AY071593 | BS01528 | FJ634634 | UGS01321 | URS01321 |
| 837 | LD34464 | AY119609 | BS10149 | FJ636575 | UGS01224 | URS01224 |
| 999 | RE02181 | AY121654 | BS11324 | FJ636822 | UGS01270 | URS01270 |
| 1011 | RE40955 | AY119633 | BS01472 | FJ634621 | UGS01303 | URS01303 |
| 1563 | GM14158 | AY058487 | BS06640 | FJ635573 | UGS01368 | URS01368 |
| 1314 | LD23745 | AY061321 | BS06812 | FJ635628 | UGS01373 | URS01373 |
| 3384 | LD43531 | BT004866 | BS10704 | KX801354 | UGS01455 | URS01455 |
| 1362 | RH33703 | AY071719 | BS13820 | KX802974 | UGS01291 | URS01291 |
| 1395 | LD26976 | AY051743 | BS08035 | FJ636068 | UGS01201 | URS01201 |
| 2112 | RE09535 | BT003533 | BS01601 | FJ634654 |  | URS01311 |
| 882 | GH03058 | AY047523 | BS07387 | FJ635837 | UGS01386 | URS01386 |
| 1140 | GH11957 | BT012317 | BS16504 | FJ638153 | UGS01419 | URS01419 |
| 2169 | GH14435 | AY051486 | BS01285 | FJ634586 | UGS01319 | URS01319 |
| 1155 | LD32727 | BT012315 | BS16432 | FJ638145 | UGS01420 | URS01420 |
| 1266 | AT14262 | AY075206 | BS14251 | KX805345 | UGS01294 | URS01294 |
| 615 | LD47783 | BT014655 | BS16517 | FJ638156 | UGS01493 | URS01493 |
| 1233 | GH23156 | AY051584 | BS07804 | FJ635992 | UGS01389 | URS01389 |
| 1344 | LD24671 | AY051708 | BS06840 | FJ635639 | UGS01375 | URS01375 |
| 1698 | SD07045 | AY069821 | ND | ~ |  |  |
| 867 | LD25202 | AY058562 | BS01052 | FJ634511 | UGS01302 | URS01302 |
| 3198 | LD15641 | AY069442 | BS04093 | KX794262 | UGS01440 | URS01440 |
| 1410 | LD31082 | AY118946 | BS03268 | FJ634709 | UGS01439 | URS01439 |
| 1863 | LD31805 | AY095046 | BS08414 | FJ636146 | UGS01250 | URS01250 |
| 5514 | LD40310 | AY095051 | ND | ~ |  |  |
| 3603 | LD33094 | BT004484 | ND | ~ |  |  |
| 1233 | LD11219 | BT021404 | BS15473 | KX800932 | UGS01409 | URS01409 |
| 735 | RE22626 | AY128470 | BS16687 | FJ638182 | UGS01482 | URS01482 |
| 1395 | RE14437 | AY071070 | BS11495 | FJ636889 | UGS01275 | URS01275 |
| 1365 | RE72705 | BT004845 | BS01529 | FJ634635 | UGS01316 | URS01316 |
| 1131 | GH16214 | AY051508 | BS07493 | FJ635888 | UGS01445 | URS01445 |
| 1668 | GH12103 | BT003617 | BS09540 | FJ636440 | UGS01448 | URS01448 |
| 3147 | GH14509 | AY075338 | BS10262 | FJ636620 | UGS01258 | URS01258 |
| 1272 | RE70055 | AY070600 | BS02838 | FJ634684 | UGS01232 | URS01232 |
| 1272 | LD22902 | AY051679 | BS07991 | FJ636052 | UGS01393 | URS01393 |
| 588 | RE65766 | AY071609 | BS13216 | FJ637392 | UGS01287 | URS01287 |
| 927 | LD26077 | AY058571 | BS08023 | FJ636063 | UGS01396 | URS01396 |
| 2538 | RE55238 | AY113499 | BS13064 | KX806112 | UGS01488 | URS01488 |
| 2502 | SD04682 | AY061595 | ND | ~ |  |  |
| 1197 | LD21748 | AY118913 | BS01111 | FJ634532 |  | URS01307 |
| 1803 | RE43755 | BT001649 | BS01577 | MN238603 | UGS01310 | URS01310 |
| 1416 | GM13220 | AF132556 | BS05308 | FJ635094 |  | URS01345 |
| 549 | GH04664 | AY069045 | BS05672 | FJ635210 |  | URS01354 |
| 951 | LD19982 | AY061280 | BS04240 | FJ634892 | UGS01334 | URS01334 |
| 1293 | GH01984 | AY047517 | BS06883 | FJ635656 | UGS01379 | URS01379 |
| 2352 | LD31802 | AY129448 | ND | ~ |  |  |
| 1704 | GH02705 | AY058273 | BS07383 | FJ635836 | UGS01244 | URS01244 |
| 1095 | LD24658 | AY058548 | BS03211 | FJ634695 | UGS01241 | URS01241 |
| 1155 | GH11591 | AY069096 | BS01103 | FJ634528 | UGS01306 | URS01306 |
| 1485 | AT28346 | BT001357 | BS12663 | FJ637258 | UGS01402 | URS01402 |
| 1707 | LD10376 | BT003278 | BS15422 | KX805222 | UGS01489 |  |
| 1053 | LD29223 | AY061369 | BS10184 | FJ636586 | UGS01227 | URS01227 |
| 2982 | RE04933 | BT015238 | BS16644 | KX800772 | UGS01481 | URS01481 |
| 1101 | GH23568 | BT016127 | BS16867 | FJ638204 | UGS01422 | URS01422 |
| 1815 | RE56844 | AY118396 | ND | ~ |  |  |
| 2265 | LD22131 | BT004489 | ND | ~ |  |  |
| 810 | GM02880 | BT003290 | ND | ~ |  |  |
| 1143 | RE62261 | BT003190 | BS15893 | KX803964 | UGS01416 | URS01416 |
| 1914 | RE12569 | AY094852 | BS11463 | KX803442 | UGS01460 | URS01460 |
| 1320 | RE69884 | BT001726 | BS15848 | KX801458 | UGS01414 | URS01414 |
| 1680 | LD36183 | BT010043 | BS10152 | KX803110 | UGS01225 | URS01225 |
| 594 | SD18370 | BT011331 | BS15670 | KX805987 | UGS01412 | URS01412 |
| 1257 | GH28523 | AY051635 | BS07875 | FJ636017 | UGS01392 | URS01392 |
| 3264 | SD05105 | BT003173 | ND | ~ |  |  |
| 1437 | LD44262 | AY122191 | BS08852 | FJ636299 | UGS01211 | URS01211 |
| 2856 | LD39519 | BT016092 | BS17648 | KX802158 | UGS01436 | URS01436 |
| 1059 | LD13640 | AY069425 | BS04669 | FJ634999 | UGS01342 | URS01342 |
| 1128 | LD43873 | AY058709 | BS08844 | FJ636295 |  | URS01210 |
| 1131 | LD44591 | AY069669 | BS10160 | FJ636578 |  | URS01484 |
| 2304 | GH03852 | AY058288 | BS07293 | FJ635807 | UGS01245 | URS01245 |
| 909 | RE29832 | AY071261 | BS12114 | FJ637073 |  | URS01280 |
| 906 | GH10637 | AY060272 | BS07539 | FJ635898 | UGS01387 | URS01387 |
| 981 | LD14639 | BT016110 | BS15495 | KX804433 | UGS01410 | URS01410 |
| 921 | RE58406 | AY113503 | BS12834 | FJ637310 | UGS01286 | URS01286 |
| 1770 | GH22686 | AY059447 | BS01815 | FJ634662 |  | URS01320 |
| 492 | AT28881 | AY089477 | BS12705 | FJ637275 | UGS01296 | URS01296 |
| 1623 | LD23505 | AY061316 | BS17604 | KX805486 | UGS01483 | URS01483 |
| 1224 | LD25827 | AY051727 | BS08117 | FJ636103 | UGS01395 | URS01395 |
| 1590 | LP01280 | AY069713 | BS09158 | FJ636363 | UGS01255 | URS01255 |
| 972 | LD37787 | AY051890 | BS17746 | KX801950 | UGS01424 | URS01424 |
| 612 | LD39986 | AY060425 | BS08757 | FJ636257 | UGS01446 | URS01446 |
| 642 | GM06568 | AY070528 | BS04282 | FJ634913 | UGS01335 | URS01335 |
| 591 | GH11193 | AY075329 | BS05820 | FJ635243 | UGS01355 | URS01355 |
| 639 | GH01619 | BT003315 | BS06407 | FJ635478 | UGS01361 | URS01361 |
| 567 | RE42508 | AY119634 | BS12457 | FJ637190 | UGS01285 | URS01285 |
| 804 | RH23273 | AY113574 | BS13489 | FJ637477 |  | URS01290 |
| 660 | LP05860 | AY060449 | BS10637 | FJ636698 | UGS01454 |  |
| 669 | RE14786 | AY071080 | BS11707 | FJ636934 | UGS01276 | URS01276 |
| 1164 | LD04613 | AY069328 | BS04004 | FJ634827 | UGS01330 | URS01330 |
| 603 | LD21953 | AY094795 | BS06671 | FJ635584 | UGS01369 | URS01369 |
| 657 | GH24702 | AY060343 | BS07832 | FJ636005 | UGS01391 | URS01391 |
| 624 | GH09086 | AY060261 | BS07047 | FJ635703 | UGS01384 | URS01384 |
| 621 | GH03685 | AY060236 | BS06991 | FJ635684 | UGS01380 | URS01380 |
| 621 | LD44762 | AY069671 | BS08961 | FJ636326 | UGS01214 | URS01214 |
| 768 | RE17845 | AY071139 | BS11759 | FJ638836 | UGS01278 | URS01278 |
| 642 | IP08727 | BT024998 | BS14580 | FJ637714 | UGS01427 | URS01427 |
| 783 | LD47384 | AY122200 | BS09126 | FJ636357 | UGS01217 | URS01217 |
| 639 | RE70650 | AY071647 | BS13079 | FJ637379 | UGS01469 | URS01469 |
| 576 | LD34217 | AY060408 | BS08459 | FJ636168 | UGS01438 | URS01438 |
| 576 | GM13874 | AY118845 | BS05469 | FJ635129 | UGS01346 | URS01346 |
| 954 | RE74715 | AY071661 | BS02841 | FJ634686 |  | URS01288 |
| 2217 | GH03557 | AY089490 | BS10234 | FJ636610 | UGS01490 | URS01490 |
| 603 | LD21679 | AY118912 | BS05602 | FJ635155 | UGS01349 | URS01349 |
| 648 | GM14354 | AY070533 | BS05574 | FJ635152 | UGS01348 | URS01348 |
| 552 | RE42418 | AY121688 | BS12554 | FJ637216 | UGS01284 | URS01284 |
| 546 | RE63021 | BT012453 | BS16093 | FJ638061 | UGS01476 | URS01476 |
| 1611 | LD36080 | AY089553 | BS09916 | FJ636505 | UGS01495 |  |
| 576 | RE36103 | AY119135 | BS12173 | FJ637087 | UGS01465 | URS01465 |
| 2913 | GH01881 | AY058264 | BS01336 | FJ634600 | UGS01313 | URS01313 |
| 576 | GH20776 | AY119536 | BS06543 | FJ635537 | UGS01364 | URS01364 |
| 1320 | SD02309 | AY050239 | BS09319 | FJ636393 | UGS01218 | URS01218 |
| 603 | LD16419 | AY061221 | BS04210 | FJ634878 | UGS01337 | URS01337 |
| 1179 | LD41588 | AY051936 | BS08802 | FJ636270 | UGS01207 | URS01207 |
| 654 | RE02339 | AY070910 | BS16737 | FJ638185 | UGS01433 | URS01433 |
| 615 | GM13948 | AY051640 | BS05471 | FJ635131 | UGS01347 | URS01347 |
| 384 | RE10554 | AY075451 | BS11434 | FJ636865 | UGS01272 | URS01272 |
| 651 | LD31286 | AY058620 | BS10439 | FJ636661 | UGS01230 | URS01230 |
| 948 | GH19261 | BT003610 | BS14948 | FJ637837 | UGS01405 |  |
| 1224 | LD40460 | AY051918 | BS08669 | FJ636229 | UGS01428 | URS01428 |
| 1248 | GM13876 | AY058486 | BS06639 | FJ635572 |  | URS01367 |
| 1542 | LD44595 | AY122193 | BS08958 | FJ636324 | UGS01213 | URS01213 |
| 1428 | SD09476 | AY052137 | BS09382 | FJ636406 | UGS01236 | URS01236 |
| 492 | LD37115* | AY051879 | BS08610 | FJ636207 | UGS01206 | URS01206 |
| 942 | HL08111 | AY060358 | BS06645 | FJ635576 | UGS01254 |  |
| 1923 | RE64552 | AY119652 | BS13207 | KX805712 | UGS01467 |  |
| 945 | GM13370* | AY060981 | BS09612 | FJ636452 | UGS01221 | URS01221 |
| 483 | IP02725 | BT023253 | BS17067 | FJ638243 | UGS01479 | URS01479 |
| 1530 | LD08669 | AY118898 | BS05002 | FJ635028 | UGS01240 | URS01240 |
| 1458 | LD15813 | BT016105 | ND | ~ |  |  |
| 1536 | RH62992 | AY071759 | BS14010 | KX805777 | UGS01470 | URS01470 |
| 651 | GH02759 | BT004505 | BS05651 | FJ635193 | UGS01352 | URS01352 |
| 2904 | LD04007 | AY089533 | BS01623 | FJ638767 | UGS01323 | URS01323 |
| 783 | GH04031 | AY118731 | BS05664 | FJ635204 | UGS01353 |  |
| 1338 | GH17420 | AY060726 | BS09464 | FJ636424 | UGS01449 | URS01449 |
| 522 | LD14743 | AY122159 | BS02803 | FJ634675 | UGS01231 | URS01231 |
| 1656 | LD30429 | AY051781 | ND | ~ |  |  |
| 3339 | LP12374 | BT010285 | BS01347 | FJ634604 | UGS01314 |  |
| 867 | RE01069 | AY069760 | BS11302 | FJ636812 | UGS01268 | URS01268 |
| 1803 | RE73180 | BT001740 | BS15854 | KX805206 | UGS01474 | URS01474 |
| 1599 | RE55465 | BT021257 | BS18055 | KX804603 | UGS01437 | URS01437 |
| 2670 | LD45081 | AY094837 | BS10614 | KX801710 | UGS01264 | URS01264 |
| 2565 | SD04959 | AY069809 | ND | ~ |  |  |
| 651 | SD04693 | AY094963 | BS10177 | FJ636584 | UGS01226 | URS01226 |
| 387 | SD05285 | BT009942 | BS17965 | KX803730 | UGS01425 | URS01425 |
| 1482 | LD46328 | AY069690 | BS08894 | FJ636315 |  | URS01216 |
| 873 | LD43943 | AY061472 | BS09946 | FJ636509 | UGS01222 | URS01222 |
| 2196 | RE65757 | BT003184 | BS15937 | KX800618 | UGS01413 | URS01413 |
| 2034 | LD42274 | AY051953 | BS08818 | FJ636277 | UGS01253 | URS01253 |
| 1446 | LD24662 | AY051707 | BS06939 | FJ635671 |  | URS01374 |
| 1134 | RE14108 | AY071059 | BS11482 | FJ636886 | UGS01274 | URS01274 |
| 732 | LD45244 | AY061489 | BS06240 | FJ635409 | UGS01359 | URS01359 |
| 1059 | LD44083 | AY069663 | BS01082 | FJ634519 | UGS01304 | URS01304 |
| 1041 | RE13621 | AY094853 | BS11472 | FJ636882 | UGS01273 | URS01273 |
| 1395 | RE17884 | AY071140 | BS01533 | FJ634637 | UGS01308 | URS01308 |
| 1755 | SD04652 | AY052097 | ND | ~ |  |  |
| 2328 | RE08173 | BT010024 | BS16278 | KX804384 |  | URS01432 |
| 1623 | LD31064 | AY051795 | BS08505 | FJ636186 | UGS01249 | URS01249 |
| 1209 | GH23271 | AY094727 | BS07805 | FJ635993 | UGS01390 | URS01390 |
| 1458 | LD20987 | AY094792 | BS04254 | FJ634901 | UGS01338 | URS01338 |
| 1425 | GH01161 | AY047501 | BS06959 | FJ635674 | UGS01376 |  |
| 3300 | LD23779 | AY118330 | BS10207 | FJ636595 |  | URS01451 |
| 801 | RE39068 | AY118391 | BS12411 | FJ637162 | UGS01283 | URS01283 |
| 804 | LD19727 | AF160941 | BS03726 | FJ634761 | UGS01327 | URS01327 |
| 462 | RE47264 | BT012459 | BS16091 | FJ638060 | UGS01475 | URS01475 |
| 2919 | LD21888 | AY069489 | BS10855 | KX800269 | UGS01459 | URS01459 |
| 948 | GH18603 | AY060735 | BS06101 | FJ635351 | UGS01356 | URS01356 |
| 831 | GH17465 | AY060729 | BS06537 | FJ635534 | UGS01363 | URS01363 |
| 1524 | LD09973 | AY069393 | BS02253 | FJ634666 | UGS01343 | URS01343 |
| 2379 | LD25754 | AY069536 | BS10816 | KX804777 | UGS01456 | URS01456 |
| 2883 | SD03117 | AY128501 | BS10843 | KX801314 | UGS01457 | URS01457 |
| 630 | GH04443 | AY060620 | BS06895 | FJ635661 | UGS01381 | URS01381 |
| 1581 | RE12101 | AY070576 | BS11456 | FJ636874 | UGS01461 | URS01461 |
| 771 | IP02648 | BT023269 | BS17164 | FJ638261 | UGS01480 |  |
| 1224 | LD41958 | AY051942 | BS08807 | FJ636272 | UGS01208 | URS01208 |
| 1227 | LD03827 | AY119574 | BS03994 | FJ634824 | UGS01329 |  |
| 1191 | LD36412 | AY058659 | BS08591 | FJ636203 |  | URS01205 |

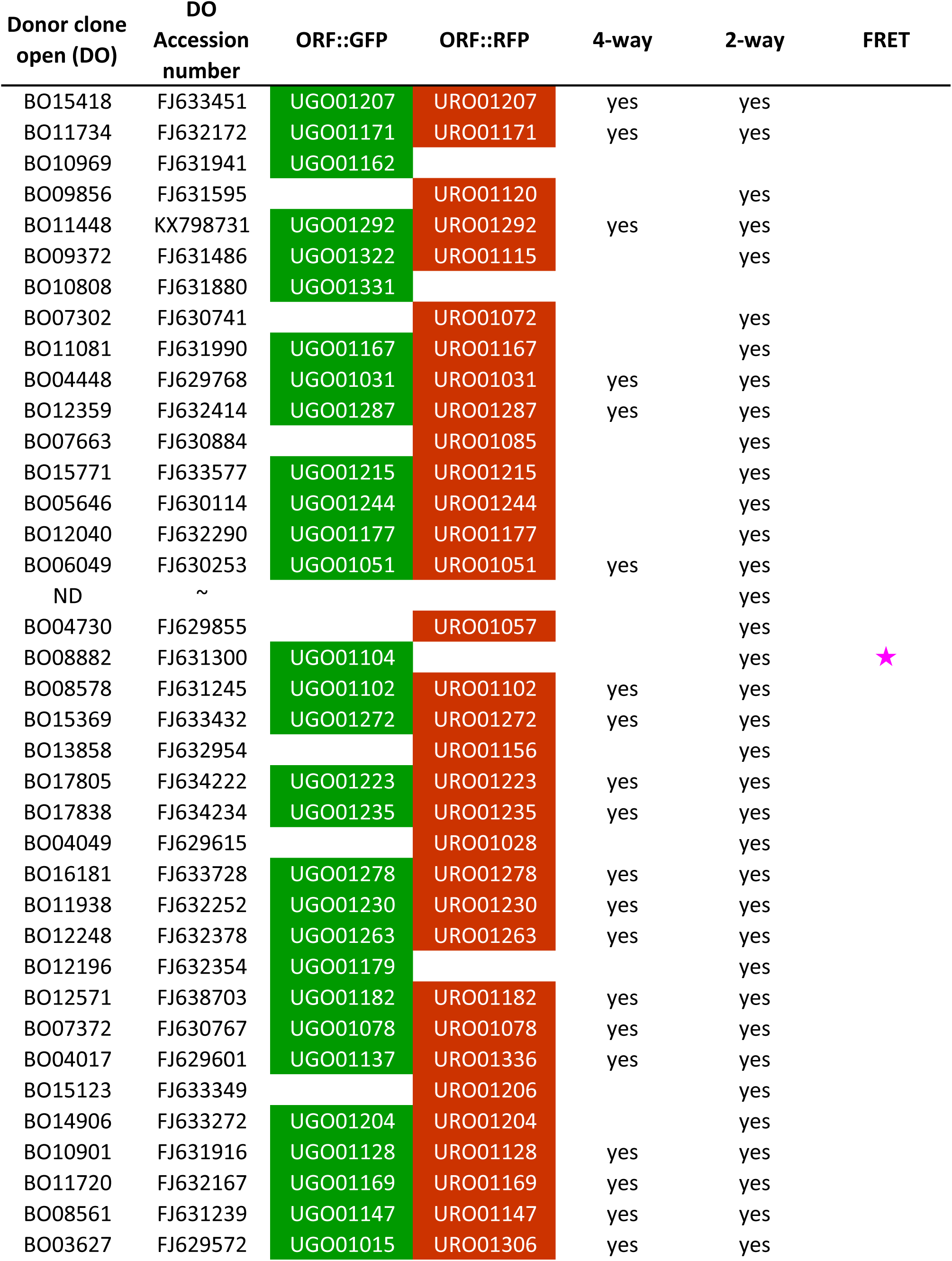
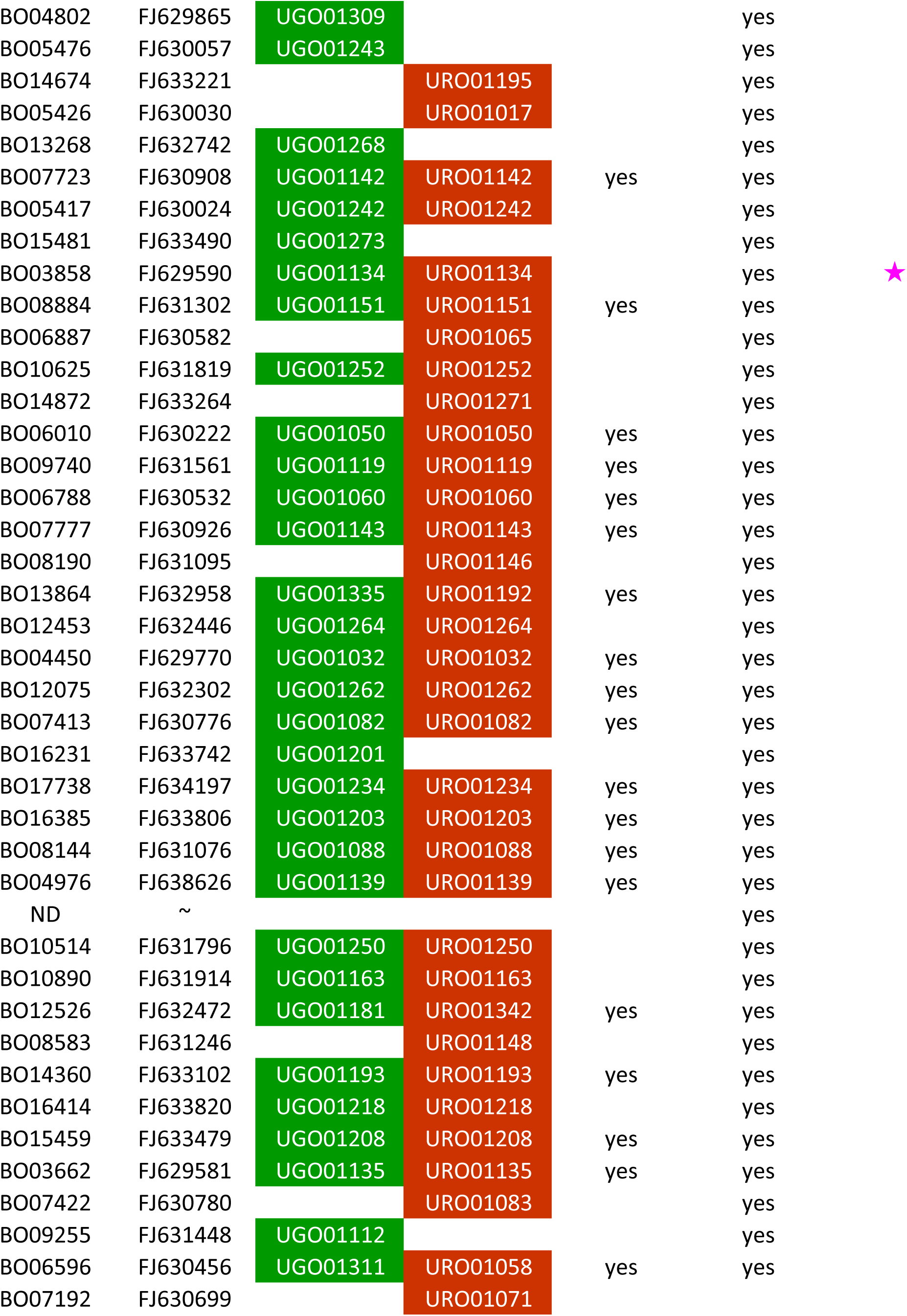
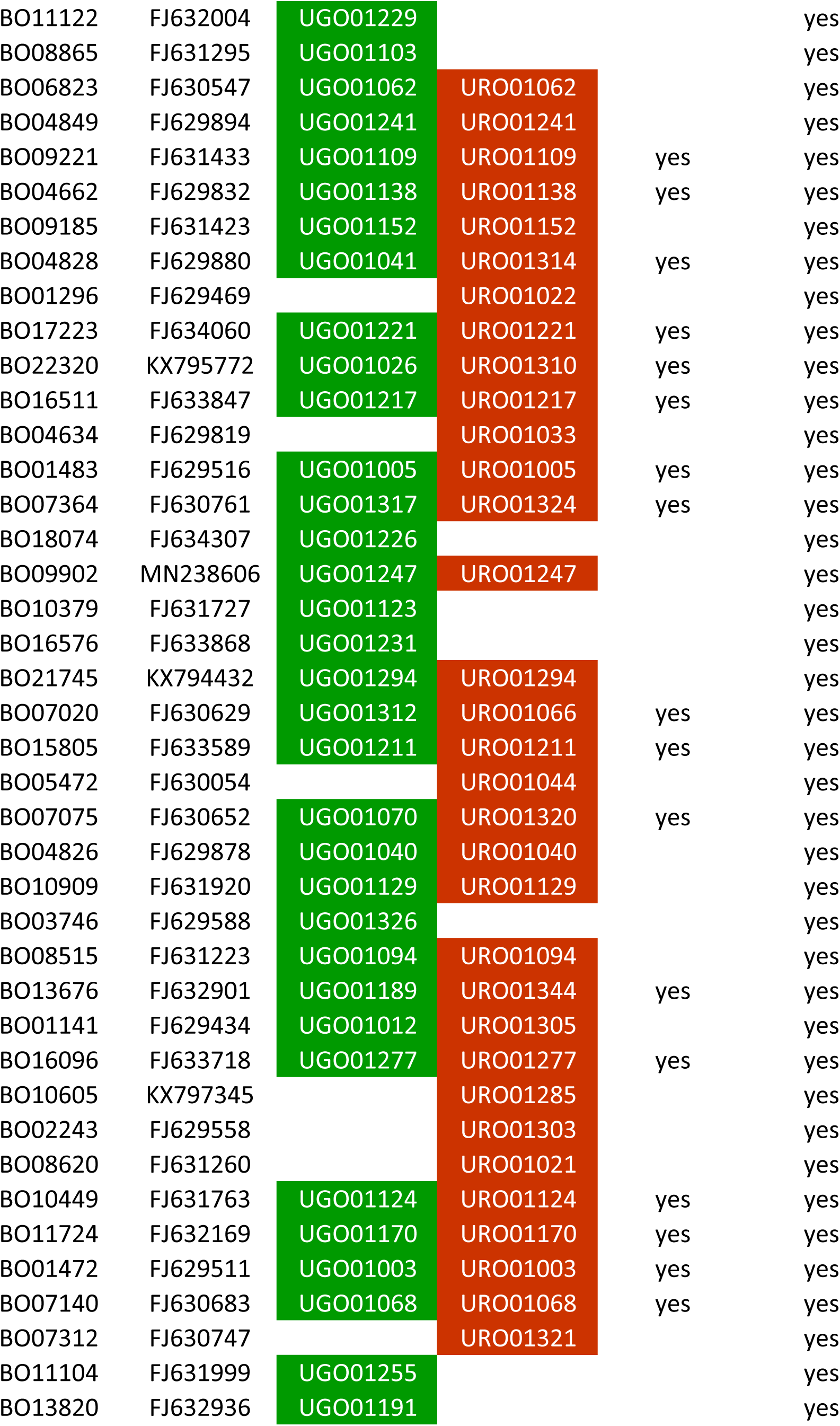
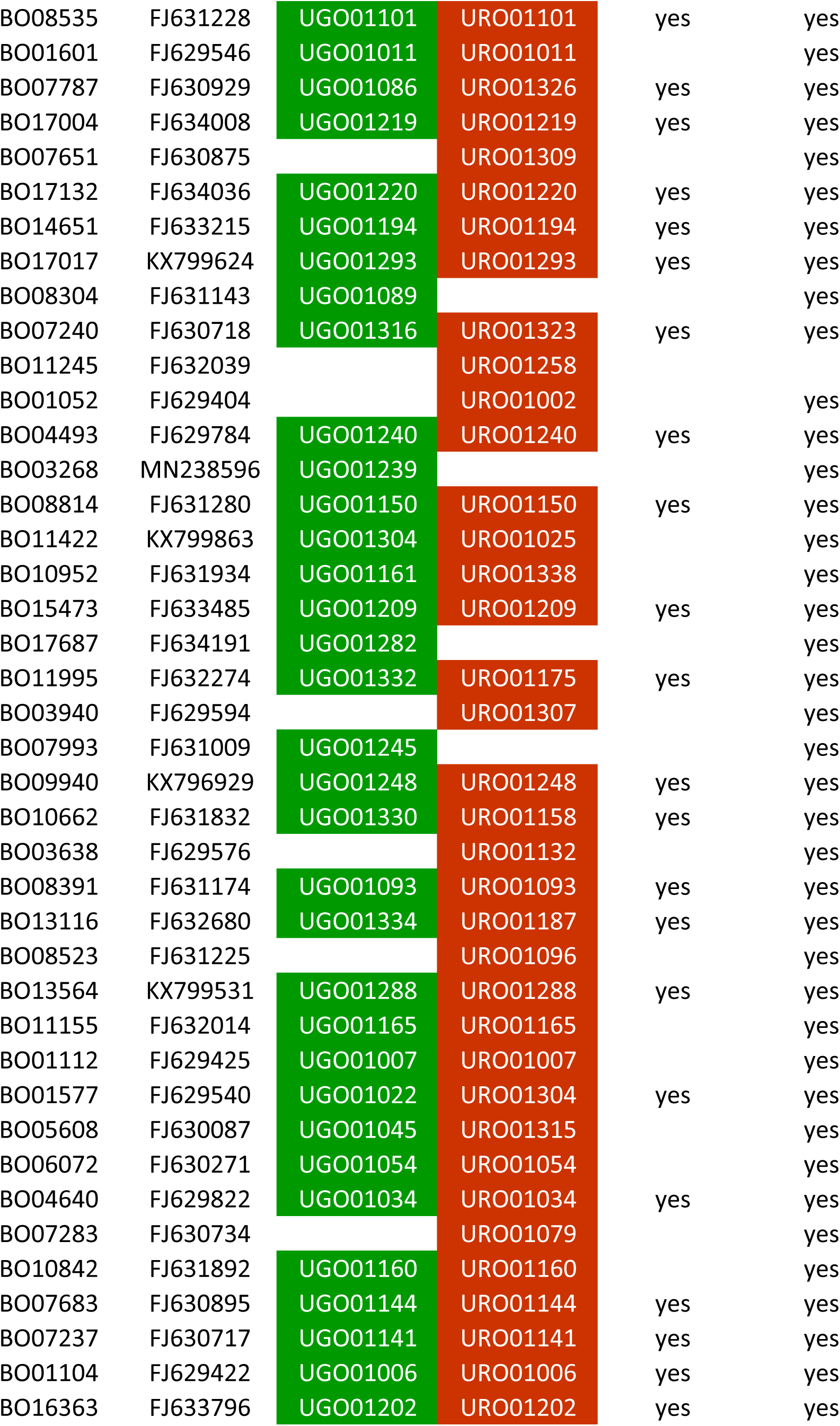
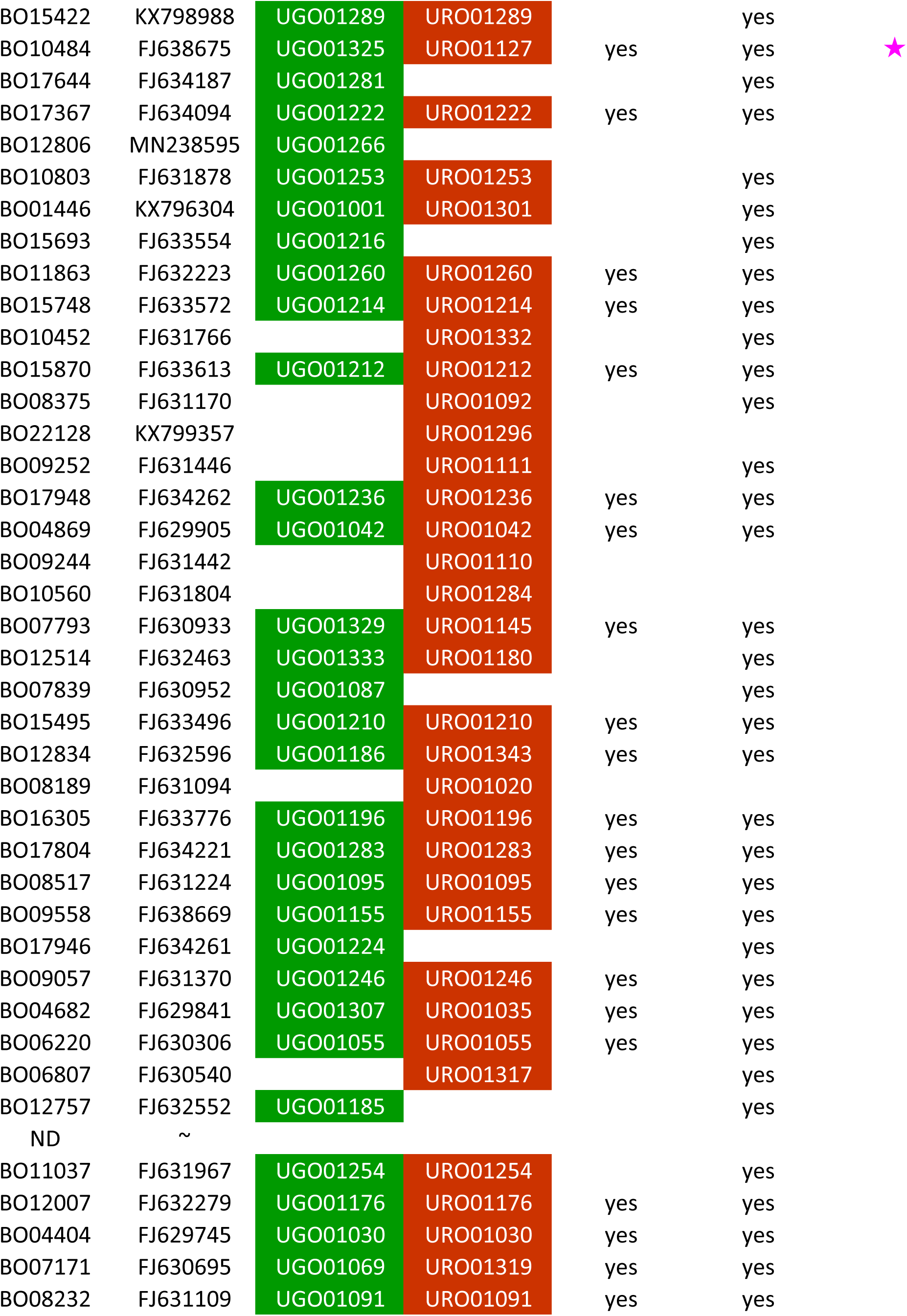
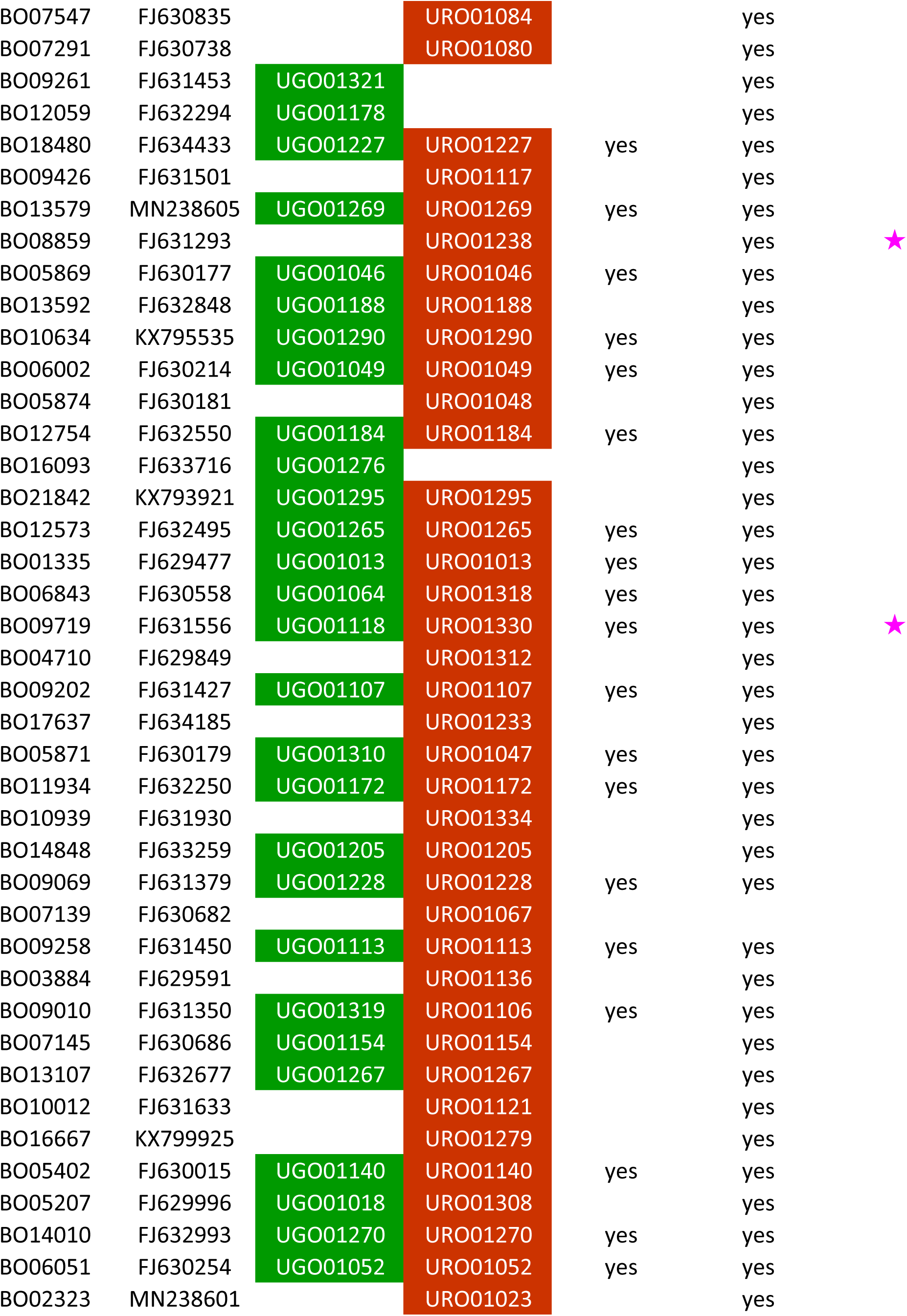
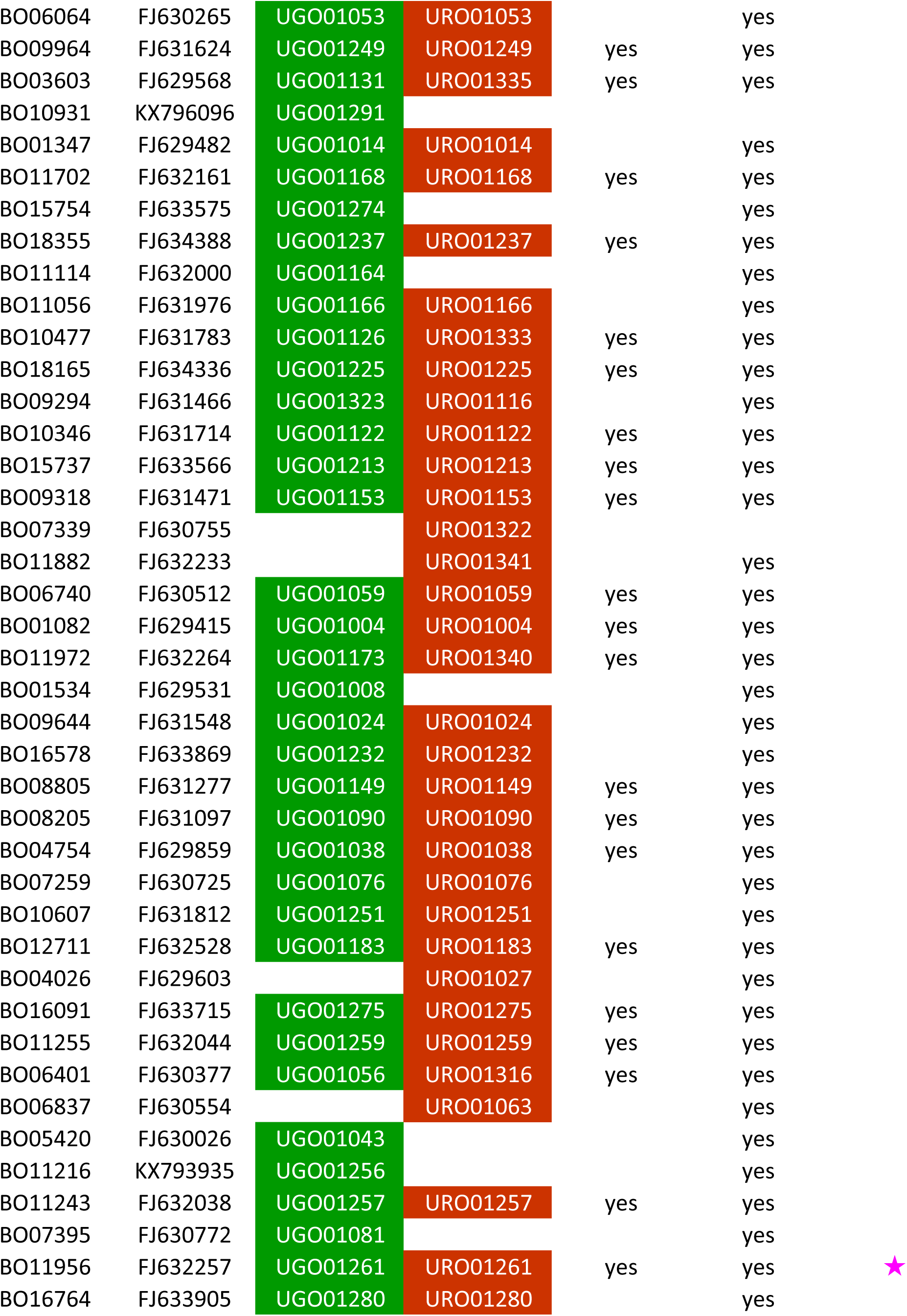
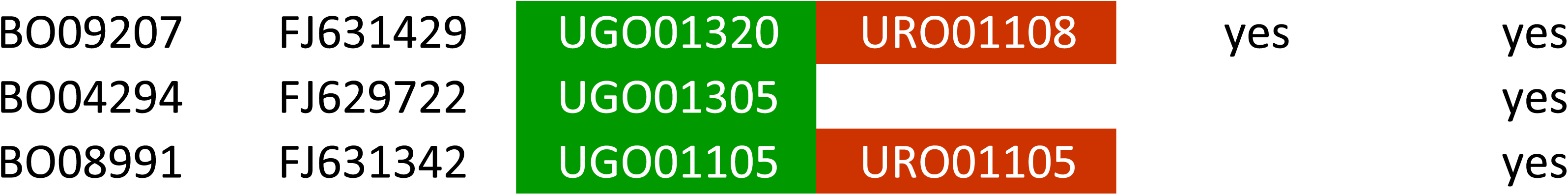

